# Gpnmb Defines a Phagocytic State of Microglia Linked to Neuronal Loss in Prion Disease

**DOI:** 10.1101/2025.02.11.637592

**Authors:** Davide Caredio, Giovanni Mariutti, Lisa Polzer, Martina Cerisoli, Beatrice Gatta, Yasmine Laimeche, Giulia Miracca, Marc Emmenegger, Marian Hruska-Plochan, Magdalini Polymenidou, Matthias Schmitz, Inga Zerr, Elena De Cecco, Adriano Aguzzi

## Abstract

Neurodegenerative conditions can induce the region-specific emergence of cell states relevant to their pathogenesis. To identify such phenomena, we generated a spatiotemporal transcriptomic atlas of mice infected with the RML prion strain. Thalamus and cerebellum experienced severe neuronal loss, developed intense microgliosis and, starting from 30 weeks post-inoculation, accumulated a novel microglial subpopulation characterized by strong expression of Glycoprotein non-metastatic melanoma protein B (Gpnmb). Elevated GPNMB levels were detected in the cerebrospinal fluid of sCJD patients, suggesting its possible usefulness as a biomarker of disease progression. The transcriptional profile of Gpnmb*^+^* microglia reflected a state of enhanced phagocytic activity with upregulation of genes associated with lysosomal function, including vacuolar ATPase V0 domain subunit d2 (*Atp6v0d2*) and Galectin-3 (*Lgals3*). In microglia-like murine BV2 cells, Gpnmb upregulation was induced by soluble find-me signals released during apoptosis, but not by apoptotic bodies or prion accumulation. Likewise, human iPSC-derived microglia showed marked upregulation of GPNMB when co-cultured with apoptotic human neurons. *Gpnmb* ablation impaired the ability of BV2 cells to clear apoptotic cells, underscoring its role in maintaining microglial phagocytosis. Our findings define *Gpnmb*⁺ microglia as a distinct, apoptosis-driven phagocytic state, linking neuronal loss to microglial activation and positioning it as a key regulator of microglial responses to prion propagation.

## Introduction

Misfolding of the cellular prion protein (PrP^C^) into aggregation-prone infectious prions (PrP^Sc^) is the causative event of prion diseases (PrDs). These pathologies are heterogeneous but inevitably lethal due to progressive neuronal death, massive glia activation (gliosis) and brain spongiosis (1). The observation that different cell types exhibit varying levels of susceptibility to infection with distinct prion strains suggests that additional factors may influence prion replication and contribute to toxicity. Some of these factors may be cell-autonomous to neurons. For example, in Creutzfeldt-Jakob disease (CJD) and Gerstmann-Sträussler-Scheinker disease, cortical and hippocampal parvalbumin-expressing interneurons show increased vulnerability and death (2, 3) whereas in fatal familial insomnia pathological hallmarks mostly occur in the thalamus without any major involvement of these neurons (4). However, the onset and the progression of motor dysfunctions correlate closely with molecular changes within glial cells and occur long before any neuronal loss is detectable (5–11). Indeed, mounting evidence highlights non-cell autonomous mechanisms in PrDs progression, suggesting that glial perturbation, rather than neuronal loss, might be the primary driver of the pathology (6, 12).

Microglia are brain resident immune cells which play a crucial role in the maintenance of CNS homeostasis and in responses to injury and disease (13). Reactive microglia are spatially associated with PrP^Sc^ in human and murine PrDs (14–17). Prion-infected, microglia-depleted mouse brains exhibit increased prion seeding activity and spread across the brain, suggesting a role for microglia in prion clearance and containment (18–20). However, microglia can also show impaired phagocytic capabilities due to overwhelming accumulation of PrP^Sc^ that cannot be effectively degraded (21). In the latter case, microglia may even facilitate the spread of prions within the CNS by migrating to different brain regions and releasing extracellular vesicles containing PrP^Sc^ (22). Additionally, oligodendrocyte precursor cells (NG2 glia) play a neuroprotective role in prion diseases by regulating microglial activity. Their selective depletion exacerbates prion-induced neurodegeneration, driven by increased prostaglandin E2 production by microglia (10). These mechanisms underscore the role of microglia not only in responding to prion infection but also in propagating it, highlighting their dual role in PrD’s pathogenesis (23).

Previous studies investigating transcriptional changes in PrDs have typically analyzed either the entire mouse brain or unique brain regions, without considering prion tropism for specific brain regions and cell types (24, 25). The detailed spatiotemporal transcriptomic analysis of PrD presented here delivers a map of the molecular landscape across various mouse brain regions during disease progression. Expression of Gpnmb identifies a distinct microglial subset emerging at 30 weeks post-prion infection and populating regions marked by intense neurodegeneration. Gpnmb is a transmembrane glycoprotein whose role in activated microglia during neurodegeneration is still unclear (26). *Gpnmb*⁺ microglia showed increased expression of the lysosomal stress marker *Lgals3* and the lysosomal proton pump subunit *Atp6v0d2,* responsible for the acidification of the luminal environment. These data suggest a role for Gpnmb in phagocytosis and lysosomal activity.

## Results

### Thalamus Emerges as the Most Vulnerable Brain Region with Profound TNF-Linked Dysregulation in Prion Disease

Eleven-week-old C57BL/6 mice were injected intraperitoneally with prion-containing brain homogenate (RML6 strain, the 6th consecutive mouse-to-mouse passage of mouse-adapted Rocky Mountain Laboratory sheep scrapie prions). Non-infectious brain homogenate (NBH) was injected as a control. At 24 weeks post-inoculation (wpi), prion-infected mice began exhibiting symptoms, including weight loss, motor impairment and increased lethargy (12). Brain samples were harvested at both 30 wpi and terminal stage of the disease (32-33 wpi, the latest timepoint at which mice could be humanely euthanized). We performed the experiment twice for each condition (prion or NBH).

Spatial transcriptomics (ST) analysis revealed a conspicuous progressive intensification of astroglial and microglial activation during the symptomatic stage of PrD. For each ST map spot, we calculated an activation score as the mean expression levels of established markers of glial reactivity (27): *Gfap*, *Vim*, *Serpina3n*, *C4b* and *B2m* for astrocytes and *Apoe*, *C1qa*, *C1qb*, *C1qc* and *Cd68* for microglia (Figure 1A). To ensure reproducibility, ST brain slices from two mice per condition were analyzed for consistency of cellular responses (Figure 1A and Supplementary Figure 1A).

**Figure 1.**
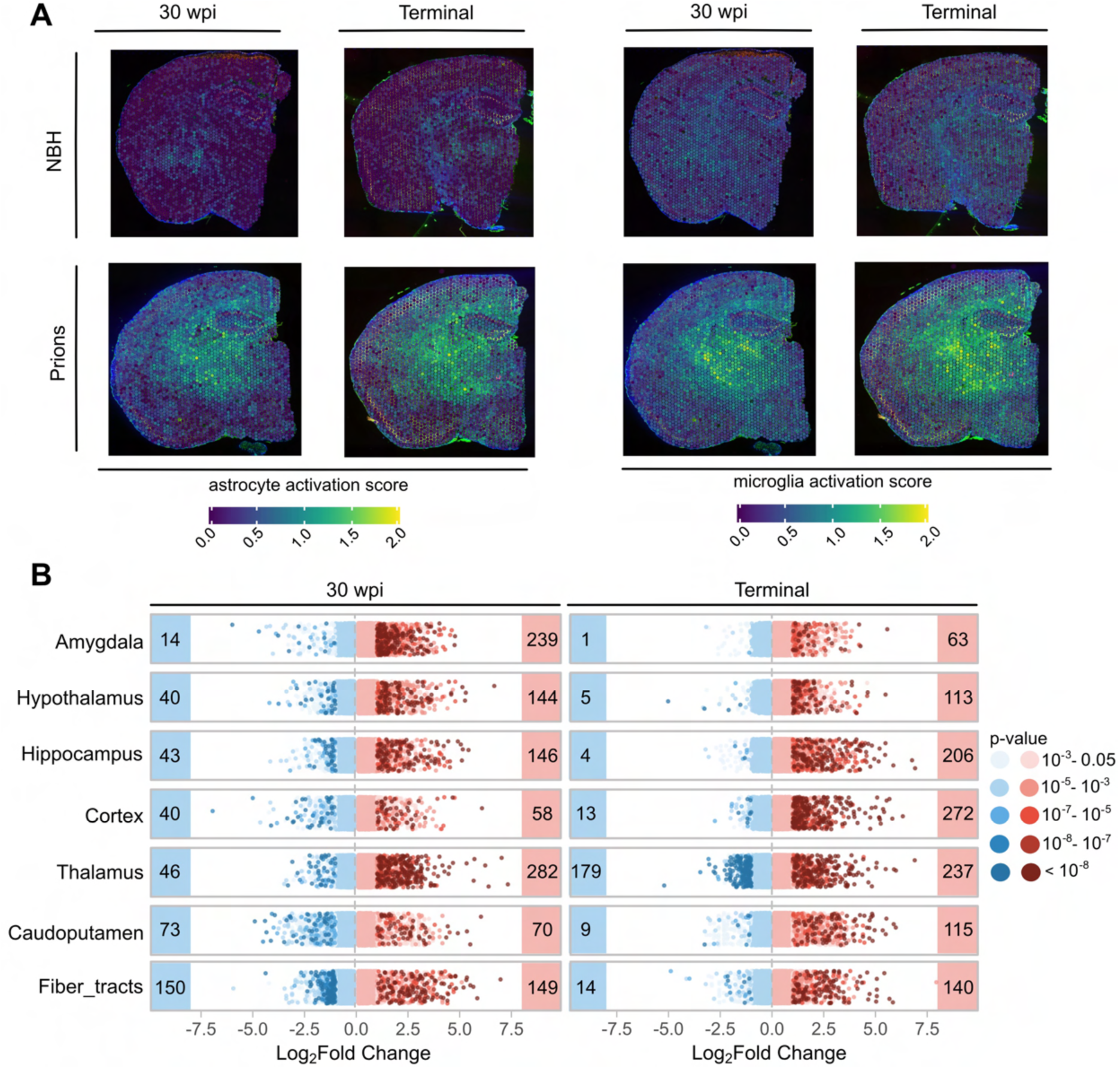
**A)** Spatial transcriptomic slices comparing prion-infected vs. uninfected mice at 30 wpi and terminal disease. Heatmaps: astrocyte (mean expression levels of *Gfap*, *Vim*, *Serpina3n*, *C4b*, *B2m*) and microglia (mean expression levels of *Apoe*, *C1qa*, *C1qb*, *C1qc*, *Cd68*) activation. **B)** Upregulated (red) and downregulated (blue) DEGs (adjusted p-value < 0.05) across brain regions at 30 wpi and terminal disease. Dots: genes with an absolute log_2_ fold change >1 color-coded according to their p-values.

Next, we investigated how distinct brain regions respond transcriptionally to prion infection. We performed differentially expressed gene (DEG) analysis between prion-infected brain regions and NBH controls (Figure 1B, and Supplementary Tables 1 and 2). At 30 wpi, we observed progressive transcriptomic deviations across multiple brain regions, including the amygdala, hypothalamus, hippocampus, and thalamus, with upregulated genes predominating (27% downregulated vs. 73% upregulated). At terminal stage, also the cortex started showing an increasing number of upregulated genes. At terminal stage the thalamus showed more downregulated than upregulated genes, whereas amygdala, hypothalamus and hippocampus maintained the same trend observed at 30 wpi.

For each filtered set of regional DEGs (|log2FC| > 0.5 and FDR < 0.05), shared modulated genes between timepoints were analyzed using over-representation analysis to identify enriched pathways (Supplementary Figure 1B and 2, and Supplementary Table 3). Most dysregulated genes across all analyzed brain regions were associated with glial cells and were related to neuroinflammatory processes, gliosis, or astrocyte development. In the hypothalamus, we detected upregulation of genes controlling cytokine regulation and leukocyte activation. In stark contrast with the rest of the brain, the thalamus displayed a unique profile of enriched immune-related pathways, with a pronounced focus on tumor necrosis factor (TNF) production, cytokine regulation and antigen presentation. This peculiar transcriptional profile, together with the extent of transcriptional dysregulation observed during disease progression, suggests that the thalamus is the brain region most affected by infection with RML6 prions.

### A distinct Gpnmb^+^ microglial subpopulation arising in late prion diseases

To identify the cell populations responsible for the observed transcriptional alterations, we applied the STdeconvolve algorithm (28) which infers gene expression profiles by analyzing the transcripts covariance and proportion of each profile at each ST pixel (Supplementary Figure 3). The algorithm then associates these profiles with cell-specific markers, hence enabling the characterization of cell-type abundance and spatial distribution across brain region during disease progression (28). At 30 wpi, no common gene expression profile emerged in two independent experiments in the prion condition (Supplementary Figure 4 and Supplementary Table 4). However, in terminally sick mice we noted the appearance of a cellular subpopulation characterized by the expression of *Gpnmb* in prion-infected samples (Supplementary Figure 5 and Supplementary Table 5). Its upregulation was already detectable at 30 wpi, particularly in the thalamus, despite not being among the most enriched genes of a subpopulation (Figure 2A and Supplementary Figure 6A). Western blots on terminally sick mice confirmed that Gpnmb was upregulated in the thalamus, hippocampus and cerebellum of prion-infected mice, but not in the hypothalamus and cortex (Figure 2B and Supplementary Figure 6B). Regions with high Gpnmb levels also showed strong PrP^Sc^ deposits (Supplementary Figure 6C).

**Figure 2.**
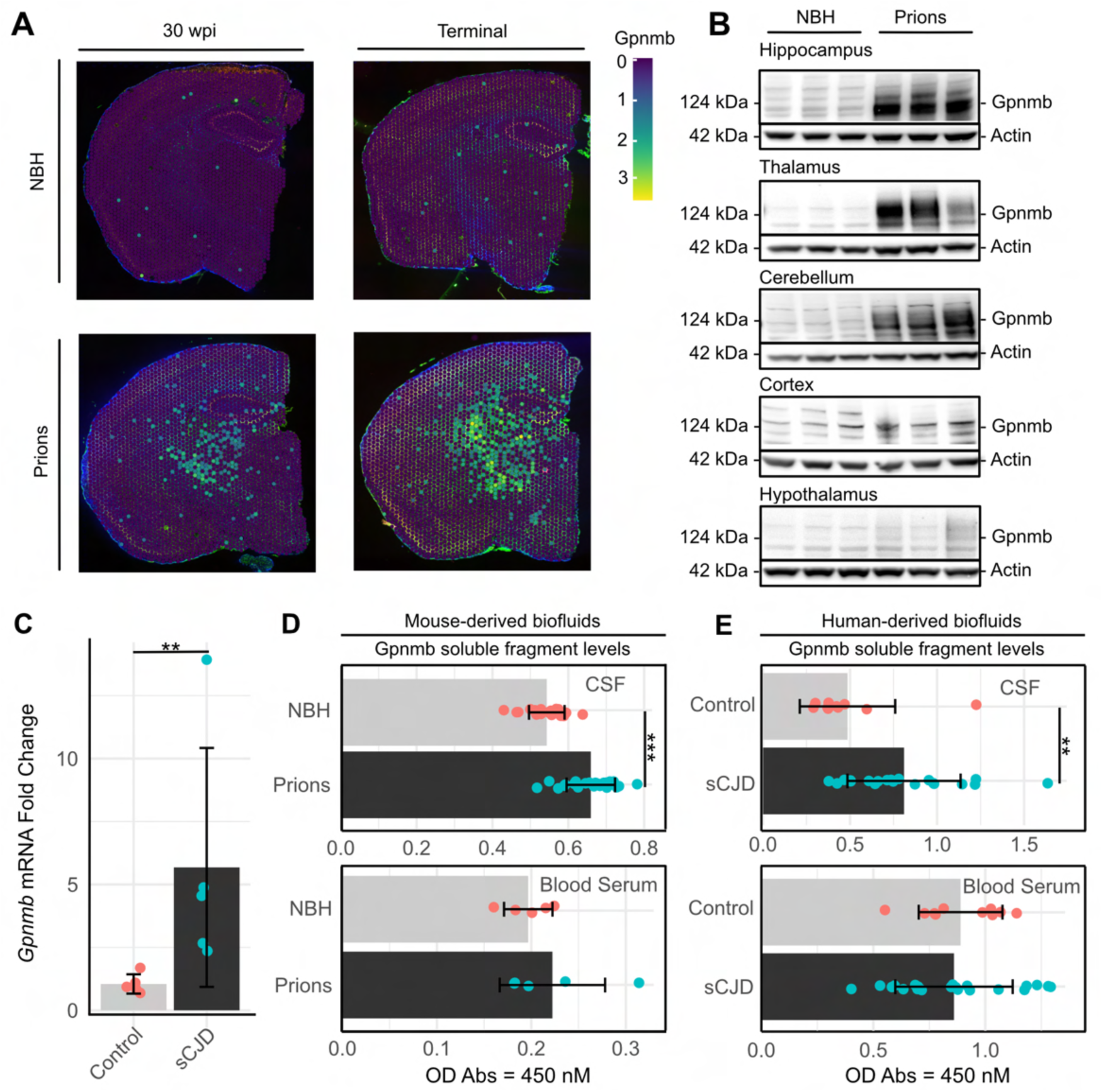
**A)** Progressive upregulation of Gpnmb mRNA in spatial transcriptomic slices of prion-infected vs. noninfected (NBH) mouse brains at 30 wpi and terminal disease. Color code: heatmap as indicated. **B)** Western blot analysis of terminally prion-sick mice displaying Gpnmb protein levels in Hippocampus, Thalamus, Cerebellum, Cortex, Hypothalamus. **C)** *GPNMB* qPCR on prefrontal cortex from sCJD and sex/aged-matched control patients. **D)** Gpnmb soluble fragment levels in cerebrospinal fluid (CSF) and blood serum from prion-infected mice and NBH controls collected at the terminal stage. **E)** Gpnmb soluble fragment levels in CSF and blood serum from sCJD patients and controls. *: p < 0.05, **: p < 0.01, ***: p < 0.005, ****: p < 0.001.

### Gpnmb and its shed fragment are increased in CJD patients

We assessed *GPNMB* mRNA levels in autopsied prefrontal cortex of five patients suffering from sporadic CJD (sCJD) and in five age and sex-matched controls (Supplementary Table 6), as this region is highly affected in sCJD (1, 29). qPCR analysis showed a significant upregulation of *GPNMB* in the prion condition (Figure 2C), underscoring the similarity of this trait in humans and mice (30).

GPNMB is known to undergo shedding by the metalloprotease ADAM10, with its soluble N-terminal domain being released in biofluids (CSF (31–35), serum (36, 37), and urine (38)). Increased shed GPNMB levels in biofluids are considered a valid biomarker of biological aging (39), but its usefulness as biomarker for neurodegenerative diseases is still debated (40–42). We observed a significant increase in shed GPNMB levels in the CSF of prion infected mice (pvalue = 4.4 × 10⁻⁷) (Figure 2D), as well as in the CSF from sCJD patients (pvalue = 0.00134871), while levels in the serum were unchanged (Figure 2E) (see Supplementary Table 7 for human cases details). These data emphasize a significant alignment in pathophysiological characteristics between human and murine PrDs.

To better understand the source of Gpnmb upregulation, we analyzed the genes co-segregating with *Gpnmb* in the molecular profiles listed by STdeconvolve (Supplementary Figure 5). The two prion-infected samples shared 19 genes belonging to the *Gpnmb*^+^ molecular profile (Figure 3A); most of them were associated with synaptic pruning and immune response, suggesting microglial involvement (Figure 3B and Supplementary Table 8). STRING pathway analysis of these genes recognized three clusters, two of which were associated with oligodendrocytes (*Mbp*, *Mobp*, *Plp1*) and microglia (*Apod*, *Apoe*, *Trf*, *Serpina3n*, *C1qc*, *C1qb*, *C1qa*, *Lyz2*, *Tyrobp*, *Ctsd*), while the third (*Gpnmb*, *Spp1*, *Lgals3*, *Vim*, *Igfbp*, *S100a6*, *Tmsb4x*) was unassigned (Supplementary Figure 7A). Therefore, we isolated and FACS sorted nuclei from major CNS cell types (microglia, neurons, oligodendrocytes and astrocytes) from prion-infected and NBH control mouse brain samples (n=3, Supplementary Figure 7B and C) and assessed Gpnmb expression levels. qPCR analysis revealed that prion-induced *Gpnmb* upregulation is driven primarily by microglia (Figure 3C).

**Figure 3.**
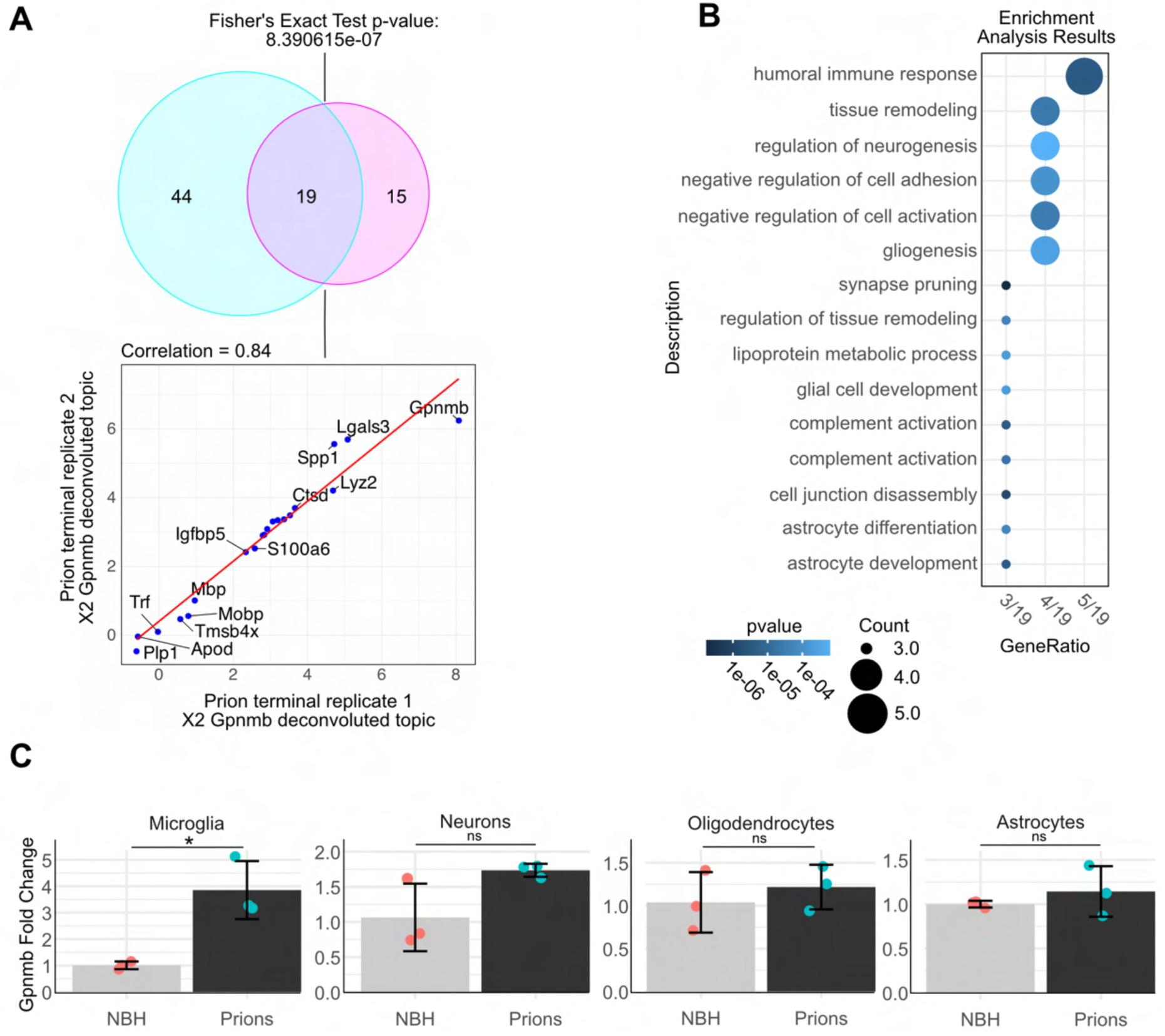
**A)** Venn diagram and correlation analysis of the ST deconvoluted Gpnmb* cell types identified in prion terminal replicates. The Venn diagram shows a significant overlap of nineteen genes (Fisher’s Exact Test, p = 8.39 × 10⁻⁷), which are displayed in the scatter plot below; the plot indicates a strong correlation (r = 0.84) between the gene expression profiles of these cell types across both replicates. **B)** Over-representation analysis (ORA) results of the nineteen overlapping genes, highlighting biological processes. **C)** qPCR showing relative mRNA abundance of *Gpnmb* in sorted CNS cell types (microglia, neurons, oligodendrocytes and astrocytes) between NBH and prion-infected samples. Statistical significance (*p < 0.05, **p < 0.01, ***p < 0.005, ****p < 0.001) is indicated by asterisks.

### Gpnmb^+^ microglia transcriptionally resembles phagocytic microglia

Microglia exist in distinct transcriptional states associated with various degrees of activation and phagocytic capacity. To better characterize the newly identified Gpnmb^+^ microglial population at the transcriptional level, we relied on a dataset of single-cell RNA sequencing of terminally ill prion-infected mice and non-infected controls (24). Unbiased subclustering analysis of the dataset (24) (Supplementary Figure 8A-C) revealed that *Gpnmb* was predominantly expressed by cells belonging to phagocytic microglia subpopulation (cluster 6) and, to a lesser extent, to MHC type-II microglia (cluster 10) only in the prion-inoculated condition (Supplementary Figure 8D and E). Among the 19 shared genes belonging to the ST *Gpnmb*^+^ profiles identified in the two prion-infected samples, only *Spp1*, *Lgals3* and *Vim* were specifically upregulated in clusters 6 and 10, whereas the others were also expressed by other microglial subclusters (Figure 4A, and Supplementary Figure 8D and E). We then analyzed the spatial expression of *Spp1*, *Lgals3* and *Vim* in our ST dataset. While *Vim* showed a broader and less specific spatial expression already at 30 wpi, *Lgals3* and *Spp1* exhibited noticeable spatial co-localization with *Gpnmb* in the thalamus and, to lesser extent, in the hippocampus at both 30 wpi and terminal stage in the prion-infected condition (Figure 4B), as confirmed also by Western Blot of prion-infected and control mouse brains (Supplementary Figure 9A and B). We hypothesized that *Gpnmb*^+^ microglia is induced by the extreme neuroinflammation characteristic of late-stage prion infections. We then conducted a targeted spatial transcriptomics experiment at 27 wpi, a stage when prion-infected mice exhibit symptomatic manifestations, including weight loss (Supplementary Figure 9C) and motor impairments, alongside ongoing microgliosis (5). Immunohistochemistry showed reactive microglia and prion deposits, particularly in the thalamus, but among the four genes, only *Vim* was upregulated at this stage (Figure 4C). This suggests that the appearance of *Gpnmb*^+^ microglial subpopulation is not a consequence of increased microgliosis.

**Figure 4.**
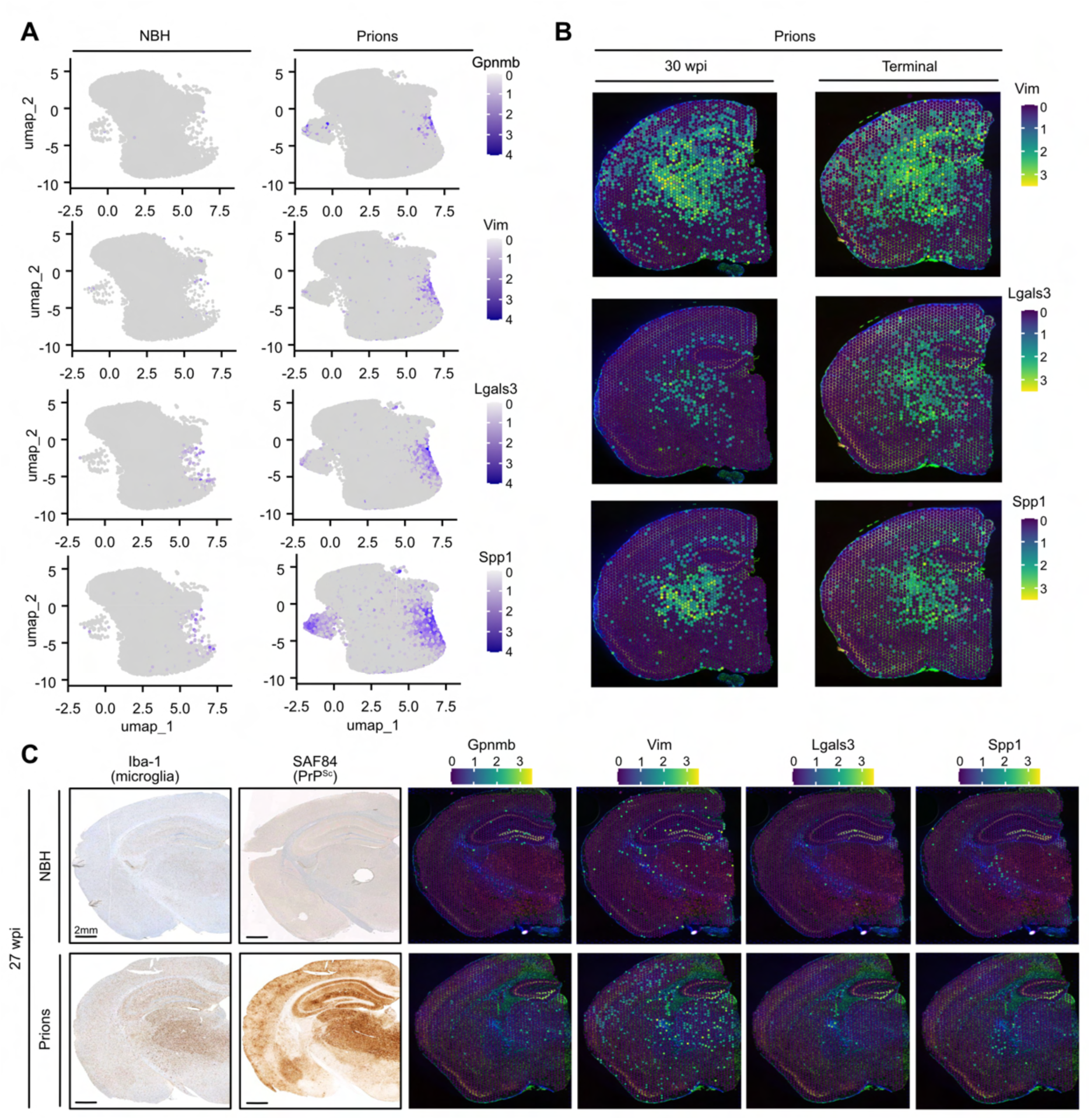
**A)** UMAP plots from a published dataset (Slota et al., 2022) displaying gene expression of *Gpnmb*, *Vim*, *Lgals3* and *Spp1* in control (NBH) and prion-infected conditions. Each plot shows the distribution of these genes within different cell populations, with higher expression indicated by darker shades. **B)** Spatial distribution of *Vim*, *Lgals3* and *Spp1* in prion-infected brain sections at 30 wpi and terminal disease. Heatmap: gene expression levels. **C)** Immunohistochemical (IHC) staining of brain sections at 27 wpi (scale bar: 2 mm) shows the microglial marker Iba-1 and PrP^Sc^ (SAF84), with spatial transcriptomics data illustrating the expression of *Gpnmb*, *Vim*, *Lgals3* and Spp1.

We conclude that the newly identified *Gpnmb*^+^ microglial subpopulation emerging during late PrD progression shares key similarities with phagocytic microglial subpopulations and is transcriptionally defined by the co-upregulation of *Lgals3* and *Spp1*. Its appearance at later stages of disease may represent a response to drastic changes in the surrounding environment and/or to unidentified secreted stimuli.

### Neuronal loss, rather than prion accumulation, triggers the appearance of Gpnmb^+^ microglia

Neuronal loss is a hallmark of late PrD that becomes widespread after the onset of cognitive and behavioral deficits (43). To test if sustained neuronal death and/or prion deposition is a trigger of *Gpnmb*^+^ microglia, we analyzed a published RNA sequencing dataset of microglial cells treated with apoptotic neurons (44). We extracted the DEGs from the phagocytic and non-phagocytic population (44), and compared it with the previously analyzed single-cell dataset on prion-infected mouse brains (24). We expressed their similarity as a Phagocytic Score. More specifically, for each microglial cell, we extracted the counts of genes associated with phagocytic and non-phagocytic conditions. The mean expression levels for phagocytic (𝜇_𝑝ℎ𝑎𝑔𝑜𝑐𝑦𝑡𝑖𝑐)_ and non-phagocytic (𝜇_𝑛𝑜𝑛−𝑝ℎ𝑎𝑔𝑜𝑐𝑦𝑡𝑖𝑐_) genes were then calculated. The Phagocytosis score’ was determined by the ratio: 𝜇_𝑝ℎ𝑎𝑔𝑜𝑐𝑦𝑡𝑖𝑐_/𝜇_𝑛𝑜𝑛−𝑝ℎ𝑎𝑔𝑜𝑐𝑦𝑡𝑖𝑐_. This ratio was visually represented as a color-coded scale to indicate the relative phagocytic activity of the microglial cells. This analysis revealed that microglia that specifically respond to neuronal loss formed a distinct subset (highlighted in purple) within broader phagocytic Cluster 6 (outlined in black) (Figure 5A). Hence microglial phagocytic responses vary depending on the nature of the engulfed material (apoptotic debris vs. prion aggregates). The subset of phagocytic microglia reacting to apoptotic stimuli overlapped with those upregulating *Gpnmb*, supporting that *Gpnmb* upregulation in microglia is driven by apoptotic neurons.

**Figure 5.**
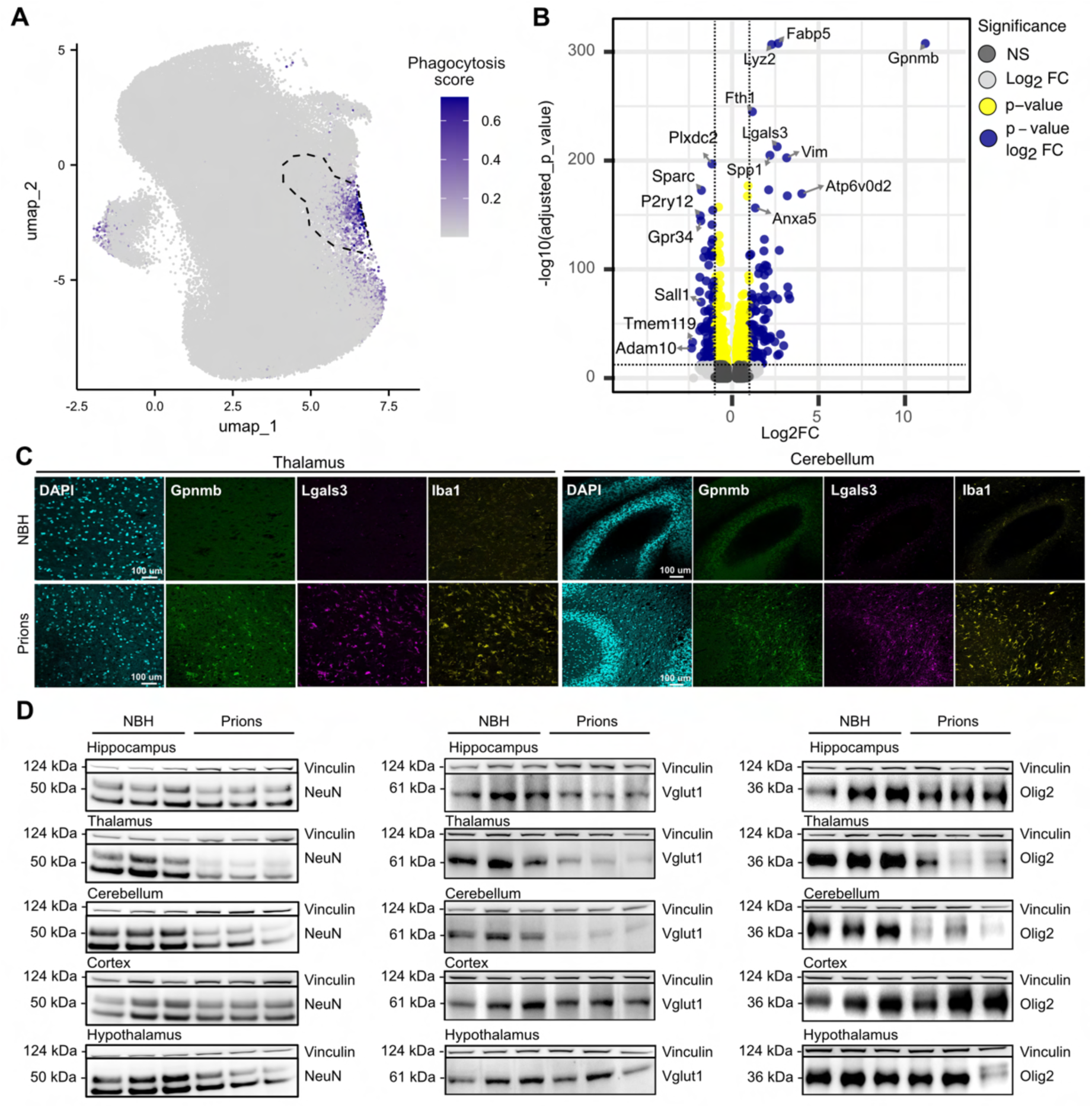
**A)** UMAP plot illustrating the distribution of the Phagocytosis score in microglial cells from single-cell RNA sequencing data, with clusters showing high phagocytic activity. Purple color coding represents higher phagocytic score expression. The dotted line represents the overall density of cluster 6. **B)** Volcano plot showing the differential gene expression analysis between *Gpnmb*⁺/Phagocytic Cluster 6⁺ and *Gpnmb*⁻/Phagocytic Cluster 6⁺ microglial cells, derived from scRNA-seq data (Slota et al. 2022). **C)** Brain sections of terminally ill prion infected mice stained with DAPI (cyan), Gpnmb (green), Lgals3 (magenta) and Iba1 (yellow). **D)** Western blot showing the levels of NeuN, a neuronal marker, Vglut1, an excitatory neuronal marker, and Olig2, mature oligodendrocytic marker, across different brain regions in controls (NBH) and prion-infected (Prions) samples at terminal disease.

We then examined gene expression in *Gpnmb*⁺ vs *Gpnmb*⁻ microglia within phagocytic Cluster 6 (Supplementary Table 9). Known phagocytosis markers were upregulated (Figure 5B), including *Anxa5* (annexin V), the receptor of phosphatidylserine, an “eat me” signal present on apoptotic bodies. *Fth1* (ferritin), which is associated with ferroptosis, a form of programmed cell death induced by iron overload during extensive phagocytosis (45), was also upregulated in *Gpnmb*^+^ microglia. The most upregulated gene in the *Gpnmb*^+^ phagocytic cluster was *Atp6v0d2*, a vesicular ATPase essential for lysosomal acidification and enzymatic activity, along with other lysosomal/phagosomal genes (Supplementary Figure 10A and Supplementary Table 9). Building on the observation that *Gpnmb*⁺ microglia in the phagocytic cluster exhibit enhanced lysosomal activity, we used Lgals3 as a marker of lysosomal dysfunction (46–48), to assess lysosomal stress in *Gpnmb*⁺ microglia. Immunofluorescence staining of terminal stage prion-infected mice revealed a robust spatial co-upregulation of Gpnmb and Lgals3 in thalamus, cerebellum and hippocampus (mostly close to the alveus and corpus callosum), but not in cortex or hypothalamus (Figure 5C, Supplementary Figure 10B-C). Although correlative, these observations indicate that exposure to apoptotic stimuli triggers the emergence of a *Gpnmb*^+^ microglial subpopulation with enhanced phagocytic activity, closely resembling that seen in prion infection. However, prion infection also leads to the presence of additional subpopulations with lower phagocytic capacity and reduced *Gpnmb* expression.

We then quantified neuronal loss in specific brain regions using the same samples shown in Figure 2B. The neuronal marker NeuN (Figure 5D, Supplementary Figure 11D) was conspicuously reduced in the thalamus, cerebellum, and hippocampus where Gpnmb, Lgals3, and Spp1 expression levels were highest (Supplementary Figures 9A and B). In contrast, NeuN levels in the cortex and hypothalamus did not show significant changes. Consistent with recent findings (43), our data indicated that the degenerating neurons were primarily excitatory (Vglut1^+^) neurons (Figure 5D middle panel and Supplementary Figure 11D), while inhibitory (Vgat^+^) neurons were largely unaffected (Supplementary Figures 11B and D). The most affected brain regions also exhibited marked synaptic degeneration (Supplementary Figure 11A and D), likely driven by the loss of Vglut1^+^ excitatory neurons. Furthermore, we observed significant oligodendrocyte loss in the cerebellum and thalamus, accompanied by extensive microgliosis in the latter (Figure 5D right panel, and Supplementary Figure 11C and D).

To identify the primary driver of Gpnmb upregulation, we treated murine BV2 cells (microglia/macrophage-like cells) with prion-infected brain homogenates, lipopolysaccharide (LPS, an inducer of general neuroinflammation), or conditioned medium from apoptotic cells. Treatment with prion-infected or NBH control brain homogenates did not modify *Gpnmb* levels (Supplementary Figure 12A). Since the total prion content in brain homogenates is stoichiometrically low, we enriched PrP^Sc^ from infected brains by NaPTA precipitation (49) (Supplementary Figure 12B). Non-infectious brain homogenates served as controls. However, even after exposure to increasing concentrations of NaPTA-purified prions (0.25%, 0.5%, and 1% w/v of the initial brain homogenate), Gpnmb upregulation was not observed (Supplementary Figure 12C). BV2 cells treated with conditioned medium from stably prion-infected but non-degenerating GT1-7 cells also failed to show any increase in Gpnmb expression (Supplementary Figure 12D). Hence, neither prion deposits nor soluble factors released during prion replication are responsible for the upregulation of Gpnmb.

To determine whether microglial Gpnmb upregulation is part of a broader inflammatory response, we treated BV2 cells with LPS, a well-known inducer of microglial activation and pro-inflammatory cytokine release. LPS exposure induced only Spp1 upregulation (Supplementary Figure 13A and B), suggesting that Spp1 plays a broader role in neuroinflammation.

To model an apoptotic environment, conditioned medium from UV-irradiated murine catecholaminergic CAD5 cells was added to BV2 cells (Supplementary Figure 13C). This resulted in a pronounced increase in Gpnmb, Spp1, Lgals3, and Atp6v0d2 expression (Figure 6A and Supplementary Figure 13D), while Vim levels remained unchanged (Supplementary Figure 13E). Notably, the robust upregulation of Gpnmb in BV2 cells was driven by soluble factors released during apoptosis, rather than by direct phagocytosis of apoptotic bodies (Supplementary Figure 14A–C).

**Figure 6.**
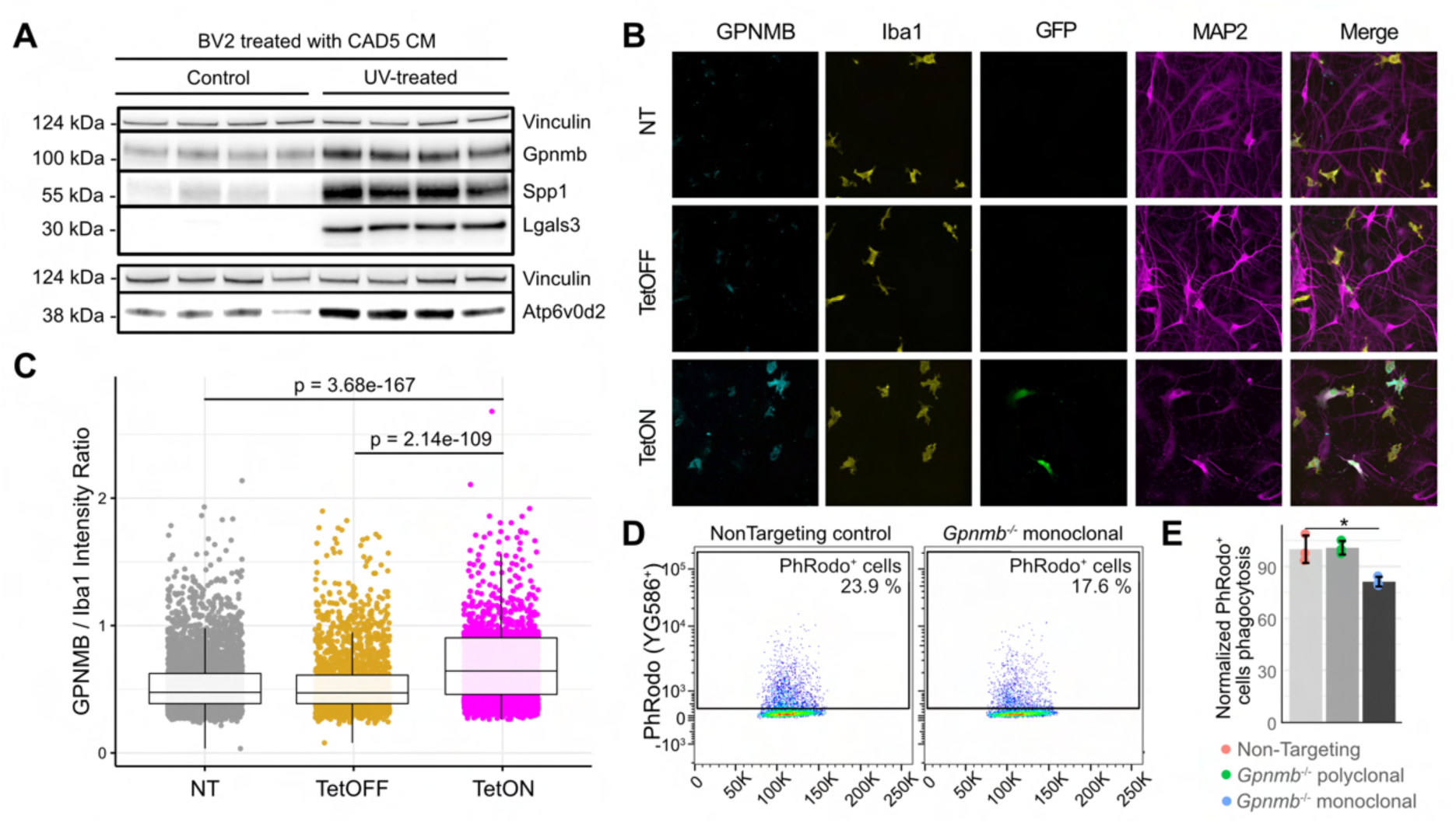
**A)** Western blot of BV2 cells treated with conditioned medium (CM) from UV-treated CAD5 cells to generate apoptotic bodies, and conditioned medium from healthy CAD5 cells for control. Gpnmb, Spp1, Lgals3 and Atp6v0d2 protein levels compared to the control protein Vinculin. **B)** Gpnmb (cyan), Iba1 (yellow, microglia marker), EGFP (green, labeling DOX-dependent expression), and MAP2 (magenta, neuronal marker). Upper row corresponds to non-treated condition (NT). Middle row corresponds to the TetOFF condition. Lower row corresponds to the TetON condition, where doxycycline is present to induce EGFP expression inducing neuronal death. **C)** Quantification of Gpnmb mean fluorescence intensity normalized to Iba1 mean fluorescence intensity across NT, DOX-OFF and DOX-ON conditions. Individual data points represent single microglia cells. Pairwise comparisons were performed using two-sided Wilcoxon rank-sum tests with Benjamini–Hochberg correction. **D)** Flow cytometry analysis of phagocytic activity in BV2 microglial cells. Cas9-expressing BV2 cells were transduced with either a non-targeting plasmid (control) or single guide RNA targeting *Gpnmb* gene, followed by selection of a monoclonal population. Cells were incubated with Phrodo-labeled apoptotic CAD5 cells. **E)** Quantification of phagocytic activity (Phrodo^+^ cells) in non-targeting control, *Gpnmb*-depleted BV2 polyclonal, and *Gpnmb*-depleted monoclonal cells (*p < 0.05).

We then utilized a human co-culture model combining iNets (induced neuronal networks) (50) with human microglia (51). iNets, derived from iCoMoNSCs (52), self-organize into synaptically connected, electrophysiologically active neuronal networks that mature into long-lived functional systems, including mixed neuronal and glial populations but lacking microglia. We then induced neuronal death by overexpressing toxic levels of EGFP (53, 54) via a doxycycline (TetON)-responsive lentiviral vector. To distinguish the effects of transgene activation, cultures were analyzed either in the presence of doxycycline (TetON), in its absence (TetOFF), or without viral transduction (NT). This enabled us to investigate the microglial response to neuronal death in a controlled environment that recapitulates human biological complexity. After 12 days of EGFP activation, we observed significant (p=1.49e-23) neuronal loss (Supplementary Figure 15A-B). In response, microglia exhibited a morphological shift from a ramified to an amoeboid shape (Supplementary Figure 15C), indicative of activation, and displayed a robust upregulation of Gpnmb (Figure 6B and C).

### Increased Gpnmb expression induces a highly phagocytic phenotype

We then directly assessed the role of Gpnmb in microglial phagocytosis. Apoptotic cells from UV-irradiated CAD5 cells were labelled with pH-Rodo-PE and added to wild-type or Gpnmb-deficient BV2 cells (Supplementary Figure 16A). Complete depletion of Gpnmb in BV2 cells significantly reduced the percentage of pHRodo^+^ cells (p < 0.05) (Figure 6D-E, Supplementary Figure 16B-D), whereas partial depletion (Supplementary Figure 16A) did not. Hence, Gpnmb enhances microglial phagocytosis, with even low expression levels being sufficient to sustain its function.

## Discussion

Microglial activation is a hallmark of prion disease, yet the molecular diversity of reactive microglial states remains poorly defined. We found that while most transcriptional changes occurred in glial cells, neuronal signals were less prominent, likely masked by extensive gliosis. The thalamus exhibited the most pronounced microglial changes, with gene ontology analysis highlighting an upregulation of TNF-related pathways, a process linked to neurodegeneration in other neurodegenerative diseases (55–58).

By deconvolving ST data into distinct cell types, we identified the emergence of a specific *Gpnmb*-expressing microglial subpopulation at the terminal stage of the disease, predominantly localized in the most affected regions. *Gpnmb* upregulation was already detectable at 30 wpi but intensified at the terminal stage, suggesting a progressive accumulation in response to sustained late-stage molecular reprogramming. Along with our finding that GPNMB and its shed soluble fragment were significantly increased in sCJD patients and prion-infected mice, these results support the notion that GPNMB upregulation is clinically relevant in PrDs of experimental animals and humans.

*Gpnmb*^+^ microglia express a distinct molecular profile including genes related to immune responses, synaptic pruning and lipoprotein metabolism as observed in other pathological contexts (59–61). Moreover, disease-associated microglial (DAM) markers such as Spp1, Lgals3 and Vim were also upregulated (44, 62, 63). Surprisingly, these changes emerged only at late disease stages, when prion deposition and gliosis were already widespread, suggesting that *Gpnmb*⁺ microglia arise in response to distinct late-stage events secondary to prion replication. As neuronal death occurs late in prion disease (43), we hypothesized that apoptotic stimuli from dying neurons might act as the primary trigger. Indeed, *in silico* analysis comparing microglia in prion condition with microglia exposed to apoptotic neurons confirmed that Gpnmb⁺ microglia belong to a transcriptionally distinct phagocytic subset specifically responding to neuronal loss. This subset not only exhibited upregulation of DAM markers previously identified with ST data (*Spp1*, *Lgals3*, *Vim*) and downregulation of homeostatic markers (*Plxdc2*, *Sparc*, *P2ry12*, *Gpr33*, *Sall1*, *Tmem119*), but also of phagocytosis-related genes including *Anxa5* (phosphatidylserine receptor), and *Fth1* (a ferroptosis-associated gene), reinforcing their role in apoptotic debris clearance. Notably, *Atp6v0d2*, a key rotor of the V-ATPase complex (64–67), was the most upregulated gene. V-ATPase has been implicated in neurodegenerative diseases (68–74) where it facilitates phagocytic digestion of apoptotic neurons (75). The upregulation of *Atp6v0d2* in *Gpnmb*⁺ microglia suggests a link to lysosomal function, as Atp6v0d2 is involved in autophagosome-lysosome fusion (76), a process essential for microglial phagocytosis. Given that Gpnmb deletion disrupts V-ATPase assembly in endothelial cells (77), its upregulation may enhance lysosomal activity by both inducing Atp6v0d2 and stabilizing the V-ATPase complex.

Supporting the hypothesis that neuronal loss triggers the emergence of *Gpnmb*⁺ microglia, we quantified neuronal density in regions showing the highest expression of Gpnmb, Lgals3, and Spp1. Strikingly, we found that NeuN, a marker for neurons, was significantly reduced in the thalamus, cerebellum, and hippocampus, regions where *Gpnmb*⁺ microglia were most abundant. In contrast, NeuN levels remained unchanged in regions such as the cortex and hypothalamus, where *Gpnmb*⁺ microglia were sparse. Further analysis revealed that the neuronal loss predominantly affected excitatory neurons, as indicated by the loss of the Vglut1 marker, while inhibitory neurons (Vgat⁺) remained largely unaffected, in line with recent observations (43). The regions experiencing the greatest neuronal loss also exhibited significant synaptic degeneration, which was likely a direct consequence of excitatory neuron depletion. In addition to neuronal loss, we observed considerable oligodendrocyte loss in the cerebellum and thalamus, particularly in the thalamus, which was accompanied by extensive microgliosis.

The emergence of *Gpnmb*⁺ microglia in prion disease as a phagocytic state responding to neuronal loss parallels other microglial populations involved in apoptotic clearance, despite not previously being recognized as a specific apoptotic-induced microglial responder. As in our immunofluorescence, its co-upregulation with Lgals3 has been observed in proliferative region-associated microglia, which engulf oligodendrocytes in developing white matter (78, 79), and in TGFβ-signaling-deficient microglia, where enhanced phagocytic activity facilitates the removal of dying Schwann cells during demyelination (79). Similarly, *Gpnmb*⁺ microglia in multiple sclerosis were found in demyelinating lesions (80). Beyond the brain, our gene ontology analysis linked *Gpnmb*⁺ microglia to rheumatoid arthritis, characterized by excessive apoptosis of osteoblasts and chondrocytes (80, 81), where Gpnmb was found to be upregulated in synovial immune cells involved in tissue remodeling (82). Comparable responses in ischemic kidney injury showed *Gpnmb*^+^ macrophages and epithelial cells were found to contain three times more apoptotic bodies compared to *Gpnmb*^-^ cells (83).

To directly test whether Gpnmb upregulation is driven by apoptotic signals (find-me cues) rather than by direct ingestion of prions (eat-me cues), we modeled apoptosis *in vitro*, given that neuronal loss in prion disease primarily occurs through autophagy or apoptosis (84). Indeed, *Gpnmb*-associated microglial markers were upregulated in murine microglia-like cells exposed to culture medium containing apoptotic bodies from UV-irradiated catecholaminergic cells, but not in cells treated with prion-infected brain homogenates or purified prions, confirming that Gpnmb induction is apoptosis-driven rather than prion-dependent. Gpnmb upregulation was still observed upon removal of apoptotic bodies from the conditioned medium, suggesting that dying cells release soluble factors that are captured by surrounding microglial cells and that trigger a drift towards a highly phagocytic *Gpnmb*^+^ state. In line with previous observation (85), Gpnmb upregulation was observed in a complex *in vitro* human system (50–52), confirming that its induction in microglia is driven by apoptotic neuronal loss rather than direct prion exposure. This further reinforces its role as a microglial responder to apoptotic stimuli, as evidenced by the morphological shift to an activated state and increased Gpnmb expression following neuronal death. The reduced phagocytic capacity in Gpnmb-deficient microglia indicates that Gpnmb facilitates apoptotic debris clearance, confirming its functional relevance. This underscores its potential role in modulating microglial responses in neurodegenerative diseases characterized by progressive neuronal loss.

Our findings establish *Gpnmb*⁺ microglia as a distinct phagocytic state in prion disease, arising in response to neuronal apoptosis rather than direct prion exposure. Their strong spatial correlation with excitatory neuronal loss, particularly in the thalamus, combined with evidence of enhanced phagocytic capacity and lysosomal activation, underscores their role in apoptotic debris clearance. Furthermore, the upregulation of GPNMB mRNA in the prefrontal cortex of sCJD patients and the increase of its shed fragment in CSF reinforce its relevance in human prion diseases, suggesting that GPNMB may serve as both a mechanistic player and a potential biomarker in prion-driven neurodegeneration. Although our *in vitro* data demonstrate that Gpnmb enhances microglial phagocytosis, these systems lack the full cellular and spatial complexity of the brain. The causal link between neuronal loss and the emergence of Gpnmb⁺ microglia remains to be firmly established *in vivo*. Future experiments inducing localized neuronal ablation, rather than inflammation alone, could help determine whether Gpnmb expression is specifically triggered by neurodegeneration and whether Gpnmb-deficient microglia exhibit impaired phagocytic function in such contexts.

## Material and methods

### Study Approvals

Animal experiments were approved by the Veterinary Office of the Canton Zurich (animal permits ZH243/2018, ZH064/2022) and carried out in compliance with the Swiss Animal Protection Law. In Switzerland, permit numbers are equivalent to approval numbers.

For the experiment in Figure 2C, prefrontal cortex cDNA from sCJD patients and age-matched controls was kindly provided by Prof. Legname’s laboratory. These samples were previously used in one of his published studies (86) under approved ethical permits. In brief, the control frontal cortex sample, obtained from an individual who died of a non-neurological condition, was generously provided by the MRC Edinburgh Brain Bank. RNA extraction was performed at the International School for Advanced Studies (Trieste, Italy), following a previously established protocol (87). Ethical approval for the use of this control sample was granted by the East of Scotland Research Ethics Service REC 1 (reference number 16/ES/0084) and informed consent for the research use of post-mortem tissue was obtained from the relatives. For the sCJD frontal cortex samples, the study was approved by the institutional review board of the Carlo Besta Neurological Institute and conducted in compliance with ethics committee guidelines. Written informed consent was obtained in accordance with the Declaration of Helsinki (1964–2008) and the Additional Protocol on the Convention of Human Rights and Biomedicine concerning Biomedical Research (2005).

All experiments and analyses involving blood samples from human donors for autoantibody screens were conducted with the approval of the local ethics committee (*Ethikkommission Kanton Zürich*, KEK-ZH-Nr. 2015-0561, BASEC-Nr. 2018-01042 and BASEC-Nr. 2020-01731) and were approved by the University of Zurich and the University Hospital Zurich, in accordance with the provisions of the Declaration of Helsinki and the Good Clinical Practice guidelines of the International Conference on Harmonisation.

The study on sCJD and related control patients (Figure 2E) conforms to the Code of Ethics of the World Medical Association. Informed consent was given by all study participants or their legal next of kin with the study being approved by the local ethics committee in Gottingen (No. 24/8/12). All samples were blinded to the investigators with respect to personal identifiers.

### Mice

Mice were kept in a conventional hygienic grade facility, constantly monitored by a sentinel program aimed at screening the presence of all bacterial, parasitic and viral pathogens listed in the Federation of European Laboratory Animal Associations (FELASA). The light/dark cycle consisted of 12/12 hr with artificial light (40 Lux in the cage) from 07:00 to 19:00 hr. The temperature in the room was 21 ± 1°C, with a relative humidity of 50 ± 5%. The air pressure was controlled at 50 Pa, with 15 complete changes of filtered air per hour (HEPA H 14 filter; Vokes-Air, Uster, Switzerland). Up to five mice were housed in IVC type II long cages with autoclaved dust-free Lignocel SELECT Premium Hygiene Einstreu (140–160 g/cage) (J. Rettenmaier and Söhne GmbH), autoclaved 20 × 21 cm paper tissues (Zellstoff), autoclaved hay and a carton refuge mouse hut as nesting material. Individual housing was avoided, and all efforts were made to prevent or minimize animal discomfort and suffering. Prion-inoculated and control-injected mice were regularly monitored for the development of clinical signs and humane termination criteria were employed.

For Visium Spatial Transcriptomics, eleven-week-old mice were intraperitoneally injected with 100 μl of RML6 prions (passage 6 of Rocky Mountain Laboratory strain mouse-adapted scrapie prions) containing 8.02 log LD_50_ of infectious units per ml. Control inoculations were performed using 100 µl of non-infectious brain homogenate (NBH) from CD-1 mice at the same dilution. Intraperitoneal injection was preferred to intracerebral injection because it allows for a slower progression of the disease and makes it possible to detect consistent neuronal loss at later stages. After inoculation, mice were initially monitored three times per week. After clinical onset, mice were monitored daily. Mice were sacrificed at pre-defined time points: 27 weeks post inoculation, 30 weeks post inoculation and terminal stage (corresponding to 31-34 weeks post inoculation). Prion-inoculated mice allocated to the terminal group were sacrificed upon clear signs of terminal PrD including piloerection, hind limb clasping, kyphosis and ataxia. Control-injected mice assigned to the latest time point group were sacrificed at the same time as terminally ill mice.

Mice were sacrificed by Isoflurane-induced deep anesthesia followed by decapitation and brains were then removed from the skull and frozen in tissue embedding medium (OCT) on dry ice.

### Spatial Transcriptomic (Visium 10X Genomics)

#### Spatial Trascriptomic Brain Slice Preparation

For RNA Quality Control and Tissue Optimization,_brain tissue from 27 wpi, 30 wpi and terminal mice undergoing ST was processed into sectioning blocks as described above. Blocks were stored in −80 °C. OCT embedded blocks were cryosectioned in a cryostat at - 20 °C to generate 10 mm slices fitting in the fiduciary frame of Visium Spatial slides, following the 10X Genomics <u>Tissue Preparation Guide</u>. A tissue optimization experiment following <u>Tissue Optimization</u> was performed with imaging of fluorescence footprint following <u>Visium</u> <u>Spatial Gene Expression Imaging Guidelines</u> on a Nikon Eclipse Ti2-E and image analysis was performed using Fiji (ImageJ).

Before undertaking a full protocol, frozen tissue was tested for RNA quality with RIN > 7.0 as shown in <u>Visium Spatial Protocols – Tissue Preparation Guide</u>, using the Agilent 4150 TapeStation. For Spatial Gene Expression, Libraries for Visium were prepared according to the <u>Visium Spatial Gene Expression Reagent Kits</u>. The quality and quantity of libraries were measured using an Agilent 4150 TapeStation. Libraries were sequenced on a NovaSeq 6000 (Illumina).

#### ST Data Analysis: Sample Preparation and FASTQ File Handling

Raw sequencing data were organized using a custom shell script. This script created symbolic links for the FASTQ files based on metadata, facilitating subsequent processing. The metadata file was parsed to link the original FASTQ files stored in a designated directory to a new working directory. FASTQ files were generated and processed using the 10x Genomics SpaceRanger (v2.0.0) software toolkit. The SpaceRanger count function was used to align the reads to the reference genome (GRCm38) and generate spatial gene expression matrices. Default parameters were used for all steps.

#### ST Data Analysis: Data Processing

We used the Seurat (v4.0.5) R package for downstream analysis of the spatial transcriptomic data. The following steps were performed: the output files from SpaceRanger, including the spatial and HDF5 files, were loaded into Seurat using custom R functions. Variable features were identified using the FindVariableFeatures function with the selection method set to “vst” and nfeatures set to 2000. The data were normalized using the NormalizeData function and scaled using the ScaleData function.

#### ST Data Analysis: Data Integration and Dimension Reduction

To integrate multiple Seurat objects and perform dimensionality reduction, we first parsed and merged the Seurat objects from previously processed data. Variable features were identified for each sample using the variance stabilizing transformation (vst) method in Seurat, selecting 2000 features for integration. The data were then normalized and scaled to prepare for dimensionality reduction. Principal Component Analysis (PCA) was conducted on the variable features and the top 30 principal components were used for further analysis. Uniform Manifold Approximation and Projection (UMAP) was then performed using the first 30 principal components to visualize the spatial transcriptomic data in a reduced dimensional space. The integration framework in Seurat was employed to identify clusters and remove batch effects from the data. Clustering was performed using the FindNeighbors and FindClusters functions. The results were visualized using DimPlot and SpatialPlot to generate UMAP plots and spatial maps, respectively. This approach allowed for detailed examination and interpretation of the spatial transcriptomic landscape, providing insights into the underlying biological processes.

#### ST Data Analysis: Astrocytic and Microglial Activation Score Visualization (*Figure 1A* and Supplementary 1A)

To assess gliosis levels, we selected a set of genes related to astrocytes (*Gfap*, *Vim*, *Serpina3n*, *C4b*, *B2m*) and microglia (*Apoe*, *C1qa*, *C1qb*, *C1qc*, *Cd68*). The expression data for these genes were extracted from each Seurat object and gliosis scores were calculated as the mean expression level of the selected genes for each cell. These scores were added to the metadata of each Seurat object. Using the svglite package, spatial feature plots were generated to visualize the gliosis scores.

#### ST Data Analysis: Differentially Expression Analysis of Different ST Regions

For the differential expression analysis, we used the MAST package (v1.12.0) to identify DEGs between different conditions and brain regions. We defined a function to perform DEG analysis for each Seurat object, comparing conditions (prion-derived and NBH control) within each brain region. We cleaned the DEG results by removing genes with patterns such as “*mt*-” and “*Bc1*” to focus on relevant genes.

### Immunohistochemistry of Brain Slices

Mouse brains were fixed in formalin followed by treatment with concentrated formic acid to inactivate prions. Brain sections (app 2μm) were generated and deparaffinized using graded alcohols followed by antigen retrieval using 10 mM citrate buffer (pH 6). The presence of prion deposits was visualized using SAF84 antibody (1:200; SPI Bio). Microglia and Astrocyte proliferation was visualized by staining the brain sections using anti Iba1 (1:2500; WAKO) and anti GFAP antibody (1:1000 Agilent technologies). Brain sections were counterstained with haematoxylin and eosin. The images were acquired using NanoZoomer scanner (Hamamatsu Photonics) and visualized using NanoZoomer digital pathology software.

### Immunofluorescence of Brain Slices

40 μm brain slices were washed with PBS thrice and permeabilized with PBS-T for 30 min. Again, three washes with PBS were performed, after which the slices were incubated in a blocking solution with 4 % donkey serum (Jackson Immuno Research, Catalog No.NC9624464) for 40 min and then in a PBS-T solution containing various primary antibodies (see below) overnight at 4 °C. The samples were washed in PBS thrice before and after staining with secondary antibodies in PBS-T for 1.5 h. Finally, they were stained with Hoechst (ThermoFisher, 62249) and mounted onto glass slides for subsequent confocal microscopy using mounting medium (Dako, S3023). All incubation steps were performed with rocking. Primary antibodies used: anti-Gpnmb (biotechne, AF2330, host goat), anti-Lgals3 (Cedarlane, CL8942AP, host: rat) and anti-Iba1 (FUJIFILM Wako, host: rabbit). Secondary antibodies: anti-goat, AF488 (ThermoFisher, A11055, host: donkey), anti-rat, AF647 (ThermoFisher, A21247, host: goat) and anti-rabbit, AF555 (ThermoFisher, A31572, host: donkey). The samples were imaged using a laser scanning inverse confocal microscope (Leica Stellaris5, with Power HyD S detectors) equipped with a supercontinuum white light laser, which was used to generate the excitation light at different wavelengths. The images in each channel were acquired using 1024 pixels in x and y and a line accumulation of 8. The stained slices were imaged using a 10-fold magnification lens (HC PL APO CS2, NA = 0.4, air) to show characteristics typical of each imaged brain region as well as with a 20-fold magnification lens (HC PL APO CS2, NA = 0.4, air) for higher resolution images. For the colocalization analysis of immunofluorescence, TIFF images containing immunofluorescence data were processed to quantify the overlap between specific protein markers (Gpnmb, Lgals3 and Iba1) across various brain regions. Each image was read and individual channels were extracted for DAPI, Gpnmb, Lgals3 and Iba1. Positive cells expressing these markers were counted by applying a threshold to each channel and a logical mask was used to identify colocalized cells. These results were combined with metadata, enabling statistical analysis. Independent t-tests were performed to compare the prion and NBH conditions across different brain regions and boxplots were created to visualize the distribution of positive cell counts between these conditions.

### Single Nuclei Isolation and FACS Sorting

Brains were extracted from RML6 infected terminal stage mice and immediately flash frozen. Half brains were used for the isolation of nuclei. Each hemisphere was incubated in 7 ml lysis buffer (11ml Nuclei PURE Lysis Buffer (Nuclei Isolation Kit, Sigma Aldrich, NUC201), 110 µl 10 % Triton X-100 (Nuclei Isolation Kit, Sigma Aldrich, NUC201), 11 µl 1 M DTT, 55 µl 40 U/µl RNAsin Plus (RNAsinPlus RNase Inhibitor, Promega, N2615)) for 10 min on ice. The tissue was then homogenized using a 30 ml dounce homogenizer and incubated on ice for 5 min. 12.6 ml of 1.8 Nuclei PURE Sucrose Mastermix (Nuclei Isolation Kit, Sigma Aldrich, NUC201) was added to the suspension, which was then resuspended and overlayed on top of 7 ml of 1.8 Nuclei PURE Sucrose Mastermix in polypropylene Beckman tubes (38.5 mL, Open-Top Thinwall Polypropylene Tube, 25 x 89 mm, Beckman Coulter, 326823). The gradient was centrifuged at 30,000 x g for 45 min at 4 °C to separate nuclei from myelin debris. The supernatant was discarded and the nuclei were resuspended in 1 ml of Nuclei PURE Storage Buffer (Nuclei Isolation Kit, Sigma Aldrich, NUC201) and transferred into a 2 % BSA in PBS (Bovine Serum Albumin, Cytiva, SH30574.02) coated FACS polypropylene tube (FalconÔ Round-Bottom Polypropylene Test Tubes With Cap, Thermo Scientific, 352063) by filtering through a 30 µm strain (MACS SmartStrainers 30 µm, Miltenyi Biotec, 130-098-462). The strainer was then washed with 1 ml of Nuclei PURE Storage Buffer four times. The suspension was then resuspended and nuclei pelleted by centrifugation at 500 g for 5 min at 4 °C. Nuclei were resuspended in 2 ml FACS buffer (2 % BSA in PBS), divided equally into two new FACS tubes and incubated for blocking with TruStrain FcX PLUS (TruStrain FcX PLUS anti-mouse CD16/32 antibody, 1:10, Biolegend, 156604) on ice for 15 min. One tube was incubated with NeuN (for neuronal nuclei, 1:100, Recombinant Alexa Fluor 488 Anti-NeuN antibody), PU.1 (for microglia nuclei, 1:50, PU.1 9G7 Rabbit mAB PE-conjugate, Cell Signaling Technologies, 81886S) and Olig2 (for oligodendrocytes nuclei, 1:2000, Recombinant Alexa Fluor 647 Anti Olig2 antibody, Abcam, ab225100) antibodies and the other tube with NeuN (same as above) and LHX2/LH2 (for astrocytes nuclei, 1:500, Anti-LHX2/LH2 antibody, Abcam, ab219983) antibodies for 45 min rotating at 4 °C. Nuclei were pelleted by centrifugation at 500 g for 5 min at 4 °C, resuspended in 1 ml of FACS buffer and counterstained with Hoechst (1:2000) for 5 min at 4 °C. Nuclei were washed twice by centrifugation at 500 g for 5 min at 4 °C and resuspension in 1 ml FACS buffer and finally resuspended in 300 µl of FACS buffer before sorting. Sorting was performed following <u>Nott, Alexi et al.</u> gating with a BD FACSAria III Cell Sorter, BD Biosciences (88).

### Single-Cell RNA Sequencing Analysis

We obtained the single-cell RNA sequencing dataset from mice, which were either prion-infected or served as control subjects (24). The data was accessed from the single cell portal at the Broad Institute through the following link: https://singlecell.broadinstitute.org/single_cell/study/SCP1962. Upon importing the raw single-cell RNA sequencing data into R, the removal of doublets and ambient RNA was performed using scrublet (89) and decontX (90), respectively. Subsequently, cells with nFeature_RNA less than 1000 or more than 7000 or mitochondrial RNA percentage exceeding 5 %, were excluded through Seurat (91). Following data normalization and regression of mitochondrial genes and cell cycle effects with Seurat, integration of data from different animals was achieved using Harmony (92). Post data integration, cell clustering was performed using UMAP and the resulting clusters were annotated based on known cell-type markers.

For the phagocytic score (Figure 5A), we utilized a previously published transcriptomic data where the authors injected apoptotic neurons into cortex and hippocampus of mice (44). Next, they sorted two microglial population: P2ry12a^+^Clec7a^-^ (non-phagocytic) and P2ry12a^-^Clec7a^+^ (phagocytic) microglia. Sorted cells underwent bulk RNA sequencing analysis to identify phagocytic vs non-phagocytic markers. The dataset was processed to calculate log2 fold change (log2FC) values, defining phagocytic markers as those with log2FC > 1.5 and homeostatic markers as those with log2FC < −1.5. To proceed, we selected a set of genes excluding those identified as unavailable in the microglial cluster from a published, prion-induced scRNAseq data (24). We extracted expression data for these genes from a subset of microglial cells. Cluster identities were appended to this data and the data was transformed into a long format for visualization. Each gene was categorized as either phagocytic or homeostatic based on the established markers. We then quantified the activity of phagocytic and non-phagocytic markers by calculating mean expression scores for each marker type within each cell cluster. These scores were integrated into the metadata of the Seurat object. For each microglial cell, the average expression of phagocytic and non-phagocytic markers was determined and matched with cluster identities, enriching the Seurat object with phagocytic_score and non-phagocytic_score. Cell IDs were extracted and matched with their corresponding cluster information. Each gene expression data point was associated with a cell ID and categorized as either phagocytic or non-phagocytic. Mean scores for each cell were calculated separately for phagocytic and homeostatic markers and added as metadata in the Seurat object. Finally, we visualized the distribution of phagocytic and non-phagocytic scores across the microglial cell population using UMAP plots. Feature plots were created to color-code cells based on their phagocytic and homeostatic scores, highlighting regions of high and low activity.

### Cell Lines Culture

BV2 and CAD5 cells were cultured in DMEM/F12 (Gibco, Thermo Fisher Scientific) supplemented with 10% heat-inactivated fetal bovine serum (FBS, Cytiva), 1% GlutaMAX (Gibco), and 1% penicillin/streptomycin (P/S, Thermo Fisher Scientific). For lentivirus packaging, HEK-293T cells were cultured in DMEM supplemented with 10% FBS. For lentiviral delivery, BV2 cells were seeded in antibiotic-free medium and placed under antibiotic selection 24 h after transduction.

For the lipopolysaccharide (LPS) treatment, BV2 cells (50’000 cells/well in 6-well plate) were administered with either 2 µg/ml LPS (E.coli O111:B4, Sigma Aldrich) or the equivalent volume of PBS (Gibco) and then incubated for 24 h. Cells were then washed once with PBS and harvested for immunoblotting.

GT1-7 cells were grown in OptiMEM (Gibco) supplemented with 10 % FBS, 1 % GlutaMAX, 1 % P/S and 1% MEM Non-Essential Amino Acids (NEAA, Gibco). Stable infection of GT1-7 cells with mouse-adapted Rocky Mountain Laboratory sheep scrapie strain prions passage 6 (RML6) was achieved as previously described (93). Briefly, cells were seeded in a 6-well plate and exposed to either 0.25 % (weight/volume) brain homogenate containing prions or to 0.25 % (w/v) non-infectious brain homogenate (NBH) as control. Cells were incubated with infectious material for 3 days, followed by a full medium change. Cells were kept in culture for at least 8 passages to ensure persistent prion replication. Stably infected GT1-7 RML and NBH cells were seeded at 70% confluency and conditioned medium was collected after 3 days from plating. BV2 cells (50’000 cells/well in 6-well plate) were treated with 1.5ml/well of GT1-7 RML and NBH conditioned medium for 48h. Cells were then washed once with PBS and harvested for immunoblotting.

### Generation of Gpnmb^-/-^ BV2 line

Genetic ablation of Gpnmb in BV2 cells was achieved using the CRISPR/Cas9 system. Briefly, BV2 cells were transduced with lentiCas9-blast plasmid (Addgene, plasmid # 52962) and, 24h later, selected with 10 µg/ml blasticidin (Thermo Fisher Scientific). BV2 cells stably expressing Cas9 were transduced with CRISPR guide RNA (gRNA) targeting mouse Gpnmb (GenScript, Gpnmb-1_pLentiGuide-Puro GenCRISPR Nickase gRNA Construct, SC1786) or non-targeting (NT) control (listed as Control_16 in (94)). 24h after transduction, cells (referred to as “Gpnmb-depleted BV2 polyclonal”) were selected with 1.0 µg/ml puromycin (Thermo Fisher Scientific). A Gpnmb^-/-^ monoclonal line (referred to as “Gpnmb-depleted BV2 monoclonal”) was isolated via limited dilution, and complete knockout was confirmed via immunoblotting.

### Lentivirus production

HEK-293T cells were seeded at 60% confluency. 24h later, cargo plasmid was co-transfected together with pCMV-VSV-G (Addgene, plasmid #8454) and psPAX2 (Addgene, plasmid #12260) plasmids using Lipofectamine 3000 transfection reagent (Invitrogen). 6h after transfection, medium was replaced with DMEM supplemented with 10% FBS and 1% Bovine Serum Albumin (Sigma-Aldrich). 3 days after transfection, supernatant was harvested, centrifuged at 1500g for 5 minutes, filtered through 0.45 µm filter (Whatman), aliquoted and stored at –80° C.

### Generation of apoptotic bodies

To generate apoptotic bodies, CAD5 cells were seeded at 60% confluency. 12h after seeding, cells were irradiated with UV light (40’000 J/m^2^ using Stratagene UV crosslinker) for 24h. Cells were then incubated at 37° C for 24h, and the apoptotic bodies-containing medium gently collected and used immediately. A twin flask of healthy CAD5 cells at matching confluency was used as control. Presence of apoptotic bodies in the medium was confirmed via flow cytometry. BV2 cells (50’000 cells/well in 6-well plate) were treated for 48h with 1.5ml/well of conditioned medium from UV-irradiated or healthy CAD5 cells. Cells were then washed once with PBS and harvested for immunoblotting. The same protocol to generate apoptotic bodies was employed for the phagocytosis assay.

Apoptotic bodies isolation protocol was adapted from Phan et al., 2018 (95). Briefly, apoptotic bodies were generated as described above. After the 24h incubation at 37° C, apoptotic bodies were mechanically detached via pipetting, and cell suspension was centrifuged at 3000g for 6 minutes. Supernatant was collected as apoptotic bodies-depleted conditioned medium (referred to as “Conditioned Medium”). The pellet was resuspended in 2 ml cold PBS and centrifuged at 200g for 6 minutes. The top 1 ml supernatant was collected as apoptotic bodies fraction (referred to as “Apoptotic bodies enrichment”) and resuspended in fresh BV2 medium. A twin flask of healthy CAD5 cells at matching confluency was used as control and underwent the same isolation protocol. Successful isolation of apoptotic bodies was confirmed via flow cytometry. BV2 cells (50’000 cells/well in 6-well plate) were treated for 48h with 1.5ml/well of conditioned medium (apoptotic bodies-depleted) and apoptotic bodies enrichment from UV-irradiated or healthy CAD5 cells. Cells were then washed once with PBS and harvested for immunoblotting.

### Prion precipitation and purification

Prions were purified from 10% (w/v) RML or NBH mouse brain homogenate as described by Wenborn et al. 2015 (49). The following reagent and materials were employed: Sodium phosphotungstic acid (NaPTA) (Sigma-Aldrich), OptiPrep density gradinet medium (Stemcell Technologies), Protease type XIV from *Streptomyces griseus* (Sigma-Aldrich), benzonase nuclease (Sigma-Aldrich), EDTA 0.5M, pH 8,0, RNA-free (Invitrogen), Sodium lauroylsarcosine (Sigma-Aldrich), Dulbecco’s phosphate-buffered saline (D-PBS) lacking Mg^2+^ and Ca^2+^ (Gibco), Ultrafree-MC microcentrifuge filtration unit (Millipore, UFC30HV00).

BV2 cells (50’000 cells/well in 6-well plate) were treated for 48h with NaPTA-precipitated RML or NBH material, amount equivalent to 0.25 %, 0.5 % or 1 % (w/v) of the initial brain homogenate in 1.5ml/well DMEM supplemented with 10% FBS and 1%GlutaMAX. Cells were then washed once with PBS and harvested for immunoblotting.

### Flow cytometry

To assess the presence of apoptotic bodies in the conditioned medium, apoptotic marker phosphatidyl serine was stained using fluorescently labelled annexin-V (BD biosciences, 556421). Briefly, 5µl of PE-annexin V were added to 100µL of conditioned medium from UV-treated and heathy CAD5 conditioned medium. After 15 minutes of incubation in the dark, samples were washed twice with PBS and resuspended in FACS buffer. Samples were acquired at LSRFortessa flow cytometer (DB biosciences). The same protocol was used to confirm apoptotic bodies isolation.

For the phagocytosis assay, UV-treated CAD5 cells were conjugated to pHrodo fluorescent dye (pHrodo red SE, Invitrogen) according to manufacturer’s protocol. Briefly, cells were detached through mechanical pipetting and resuspended at 10^6^ cells/ml in PBS. Substrates were incubated in 1mM pHrodo for 2h at room temperature in the dark. After conjugation, substrates were washed 3 times in PBS and used immediately. BV2 cells (100’000 cells/well in 24-well plate) were treated with 100µl pHrodo-labelled CAD5 apoptotic bodies (around 100’000 apoptotic cells) and incubated for 1.5h and 3h at 37°C. After incubation, samples were washed twice with PBS and resuspended in FACS buffer. 10’000 events were acquired per sample using LSRFortessa flow cytometer (DB biosciences). Phagocytosis was quantified as percentage of pHrodo^+^ cells compared to untreated sample. The experiment was performed 3 times, each one in 3 technical replicates. Each dot in the graph represents the average of 3 technical replicates. All flow cytometry data were analyzed using FlowJo 10 (Tree Star).

### Immunoblotting and silver staining

Mouse samples were homogenized twice at 5,000 rpm for 15s in 1 ml of cell-lysis buffer (20 mM Hepes-KOH, pH 7.4, 150 mM KCl, 5 mM MgCl_2_, 1 % IGEPAL) supplemented with cOmplete mini protease inhibitors (Roche) using a Precellys24 Sample Homogenizer (LABGENE Scientific SA, BER300P24). After 20 min incubation in ice cleared lysates were obtained by centrifugation at 2,000 x g, 4° C for 10 minutes.

Cell extracts and NaPTA-precipitated samples were prepared in lysis buffer (50 mM Tris–HCl pH 8, 150 mM NaCl, 0.5 % sodium deoxycholate and 0.5 % Triton-X 100) supplemented with cOmplete mini protease inhibitors (Roche). In case of proteinase K (PK) (Roche) digestion, proteinase inhibitors were avoided.

Total protein concentration was measured using bicinchoninic acid assay (BCA) according to the manufacturer’s protocol (Pierce). Western blots were performed using standard procedures. Briefly, samples were boiled at 95 °C in 1X NuPAGE LDS sample buffer (Invitrogen) supplemented with 1 mM DTT (Sigma-Aldrich) loaded into NuPAGE Bis-Tris precast PAGE gels (Invitrogen) and transferred on PVDF membranes (Invitrogen).

For NaPTA-precipitated prions, Proteinase K digestion was performed at 10 µg/ml for 1h at 37° C. For silver staining, loading of NaPTA-precipitated samples was equivalent to 100µl of 10% (w/v) of the initial brain homogenate. As control, 2µl of 10% (w/v) normal NBH and RML brain homogenate were loaded. Silver staining was performed using SilverXpress kit according to manufacturer’s instructions (Invitrogen).

Following are the antibodies used for immunoblotting and their working dilution: anti-GPNMB 1:1000 (Bio-Techne, AF2330), anti-LGALS3 1:2000 (Abcam, ab2785), anti-SPP1 1:3000 (Abcam, ab11503), anti-VIM 1:3000 (Abcam, ab92547), anti-ATP6V0D2 1:2000 (Novus Biologicals, NBP3-10978), anti-NeuN 1:1000 (Abcam, ab177487), anti-VGLUT1 1:1000 (Abcam, ab77822), anti-VGAT 1:500 (Santa Cruz Biotechnology, sc-393373), anti-Olig2 1:1000 (Abcam, ab109186), anti-Iba1 1:1000 (FujiFilm WAKO, 016-20001), anti-GFAP 1:10’000 (Dako, Z0334), anti-SYP 1:1000 (BD Biosciences, #611880), anti-PrP POM1 300ng/ml (96), anti-Vinculin 1:5000 (Abcam, ab129002), anti-Actin-HRP 1:10’000 (Sigma, A3854), anti-Rabbit_IgG-HRP 1:10’000 (Jackson ImmunoResearch, 111.035.045), anti-Mouse_IgG-HRP 1:10’000 (Jackson ImmunoResearch, 115.035.003), anti-Goat_IgG-HRP 1:10’000 (Jackson ImmunoResearch, 705.035.147), anti-mouse_IgM-HRP 1:1000 (Zymed, 61-6420).

### qPCR sample preparation

CJD bulk tissues were lysed in TE buffer with the anionic detergent sodium dodecyl sulphate (SDS) and digested at 50 °C with 2 mg ml^-1^ Proteinase K (PK) for 2 h to eliminate solids and release DNA/RNA from proteins. Although prions are well-known for their relative resistance to PK digestion, prion infectivity largely depends on PK-sensitive oligomers. Indeed, prolonged PK digestion reduces prion titers by a factor of > 10^6^, but residual PK-resistant material may still be infectious. In a second step, TRIzol (TRIzol LS Reagent, Invitrogen, 10296028) reagent solution was added to the lysate (it contains Gdn-SCN and phenol, which inactivate RNases and disaggregate prions) and kept overnight at 4 °C. Gdn-SCN is a chaotropic salt which rapidly denatures proteins and abolishes the infectivity of PrD inoculum. At high concentrations, guanidine salts disaggregate PK-resistant PrP^Sc^ fibrils, eliminate PK resistance and abolish PrP^Sc^ conversion, meaning that any PK-resistant material that survived the digestion step would be expected to be inactivated at this stage of the protocol. 0.2 ml of ultrapure phenol:chloroform:isoamyl alcohol (Thermo Fischer Scientific) was added to the samples, followed by strong shaking and incubation at room temperature for 5 mins. Centrifugation step at 12 g for 15 min at 4 °C generated two phases. The aqueous upper phase was transferred to a fresh tube; 0.5 ml of isopropanol and 1 µl of Glycoblue Coprecipitant (Thermo Fisher Scientific) were added. Next, RNA was pelleted for 20 min at 12 g at 4 °C and washed twice with 75 % ethanol. The RNA pellet was dissolved at 55 °C in 11 µl of free nuclease water.

For mouse samples, the tissue was homogenized in TRIzol for 5 min at 50 oscillations/s using a Precellys24 Sample Homogenizer (LABGENE Scientific SA, BER300P24). Samples were left in TRIzol overnight at 4 °C. Samples were then warmed up to room temperature and 400 μl chloroform was added, followed by vigorous mixing. The mix was incubated for 2 min at room temperature and centrifuged at 16 g for 30 min at 4 °C. The upper phase was transferred in a new tube, 0.5 µl GlycoBlue Coprecipitant Transfer the aqueous upper phase to a new tube, 0.5 µl GlycoBlue Coprecipitant (Thermo Fisher Scientific) and 500 µl isopropanol were added, followed by vortexing. The mix was incubated for 10 min at room temperature and the RNA pelleted by centrifuging at 16 g for 30 min at 4 °C. The supernatant was discarded and the pellet washed twice with 75 % ethanol by centrifuging at 16 g for 10 min at 4 °C. The RNA was then dissolved in 11 µl RNase-free water for 10 min at 65 °C.

For BV2 cells, 50’000 cells were seeded in each well of a 6-well plate and treated the day after with 0.5 % or 1 % (w/v) NBH and RML mouse brain homogenate. After 48h incubation, BV2 cells were washed once with PBS and then harvested for qPCR. RNA was extracted using RNeasy kit according to manufacturer’s protocol (Qiagen).

RNA concentration was assessed using a NanoDrop spectrophotometer (Thermo Fisher Scientific). Reverse transcription was performed following the manufacturer’s instructions using the Quantitect Reverse Transcription kit (Qiagen). 10 ng of cDNA per sample was transferred into 384-well PCR plates (Life Systems Design) in triplicates and the detection was conducted using SYBR green (Roche) mastermix. Readout was executed with ViiA7 Real Time PCR systems (Thermo Fisher Scientific). qRT-PCR data was analyzed using the 2^-ΔΔCT^ method and visualized using GraphPad Prism.

Primer sequences: mouse GPNMB (fw: 5’-TCT GAA CCG AGC CCT GAC ATC-3’, rev: 5’-AGC AGT AGC GGC CAT GTG AAG-3’), GPNMB human (fw: 5’-ACT GTT GCT CTT GGT GGA CG-3’, rev: 5’-CCA GGA GCA GAA ATC CCA GG-3’), GAPDH mouse (fw: 5’-CCA CCC CAG CAA GGA GAC-3’, rev: 5’-GAA ATT GTG AGG GAG ATC CT-3’), GAPDH human (fw: 5’-TGC ACC ACC AAC TGC TTA GC-3’, rev: 5’-GGC ATG GAC TGT GGT CAT CAG-3’).

### ELISA

For Gpnmb ELISA measurements in murine samples (Figure 2D), to assess samples for the binding of IgGs to GPNMB (#GPB-H5229, Acro Biosystems), high-binding 1536-well plates (Perkin Elmer, SpectraPlate 1536 HB) were utilized. These plates were coated with 1 μg/ml antigen in PBS at 37 °C for 1 h, followed by three washes with PBS 0.1 % Tween-20 and blocking with 5 % milk in PBS 0.1 % Tween-20 for 1 h. 3 μl plasma, diluted in 57 μl sample buffer (1 % milk in PBS-T), were dispensed at various volumes into the antigen-coated 1536-well plates using contactless dispensing with an ECHO 555 Acoustic Dispenser (Labcyte). Dilution curves ranging from plasma dilutions 1:50 to 1:6000 were generated, with eight dilution points per patient plasma sample. The chosen ECHO liquid calibration for this assay (1:20 pre-diluted inactivated plasma sample in PBS 0.1 % Tween20) was SP2, accounting for a water-based solution containing proteins and detergents affecting the liquid’s surface tension. Following a 2 h sample incubation at room temperature, the wells were washed five times with wash buffer and the presence of IgGs bound to the antigen was detected using an HRP-linked anti-human IgG antibody (Peroxidase AffiniPure Goat Anti-Human IgG, Fcγ Fragment Specific, Jackson, 109-035-098, at 1:4000 dilution in sample buffer). Incubation of the secondary antibody for one hour at room temperature was followed by three washes with PBS 0.1 % Tween-20, addition of TMB, a three-minute incubation at room temperature and the addition of 0.5 M sulfuric acid. The well volume for each step reached a final volume of 3 μl. Plates were centrifuged after all dispensing steps, except for the addition of TMB. Absorbance at 450 nm was measured using a plate reader (Perkin Elmer, EnVision).

For GPNMB ELISA measurements in human samples (Figure 2E), high-binding 384-well plates (Perkin Elmer, SpectraPlate 384 HB) were coated with 1 μg/mL of anti-human Osteoactivin/GPNMB antibody (Novus Biologicals, AF2550) in PBS. After overnight incubation at 4 °C, the plates were washed three times with PBS-T and blocked with 5 % SureBlock (LubioScience, SB232010) in PBS-T for 1 h at RT. The sCJD CSF and serum samples, controls and the standard (R&D Systems, 2330-AC) were diluted with 1 % SureBlock (PBS-T) and transferred to the coated plates. After 2 h of incubation at RT, the plates were washed five times with PBS-T and 1 μg/mL of biotinylated anti human Osteoactivin/GPNMB antibody (Novus Biologicals, BAF2550) diluted in 1 % SureBlock (PBS-T) was added. Following an incubation of 1 h at RT, plates were washed three times with PBS-T and Streptavidin-HRP (Biologend, 405210) diluted 1:1000 in 1 % SureBlock (PBS-T) was added for 30 min at RT. After three washes with PBS-T, TMB (Invitrogen, 10354603) was added for color development and the reaction was stopped after 5 min with 0.5 M sulfuric acid. The absorbance at 450 nm was measured using a plate reader (BMG Labtech, FLUOstar Omega). CSF samples were diluted 1:20, while serum samples were diluted 1:40. All samples were run in triplicates.

### iNets generation and subculture

iNets (50) were differentiated from iPSC-derived self-renewing human neural stem cell line (iCoMoNSCs) (52) obtained from control human skin fibroblast, as described previously (50). In brief, 600,000 iCoMoNSCs were plated onto Matrigel-coated 6-well plates (Corning, 354234) in NSC medium - DMEM/F12 medium (Gibco, 11330020); 0.5× B27– supplement (Gibco, 12587-010); 0.5× N2 supplement; 1× GlutaMAX (Gibco, 35050-061) and 25 ng/ml bFGF (Gibco, PHG0261) - and complete medium was changed daily until cells reached ∼95% confluency. At this stage, NSC medium was switched to D3 differentiation medium - DMEM/F12 (Gibco 11330032); 0.5× B27+ supplement (Gibco,17504-044), 1× N2 supplement (Gibco, 17502-048); 1× GlutaMAX (Gibco, 35050-061); 1× Penicillin/Streptomycin (Sigma, P4333-100ML) - supplemented with 5 μM Forskolin (Cayman, AG-CN2-0089-M050), 1 μM synthetic retinoid Ec23 (Amsbio, AMS.SRP002-2), 500 nM Smoothened agonist SAG (Millipore, 5666600) for a total of 5 days. On the days 6–10, Ec23 was increased to 2 μM. On days 11–25, Ec23 was decreased to 10 ng/ml, SAG to 50 nM and 20 ng/ml BDNF (PeproTech, 450-02), 20 ng/ml GDNF (PeproTech, 450-10) and 20 ng/ml CNTF (Alomone labs, C-240) were added. At day 26 and onwards, medium was switched to maturation medium - 1:1 DMEM/F12:Neurobasal (Gibco, 21103049) mix; 1× B27+ supplement, 1× N2 supplement; 1.5× GlutaMAX, 5 μM forskolin, BDNF, GDNF, CNTF, NT-3 (PeproTech, 450-03) and IGF-1 (Stem Cell, 78022) all at 20 ng ml−1 and 10 μM cAMP (Sigma-Aldrich, D0260). One-third of medium was changed daily at days 0–10, whereas from this point on only two-thirds of the medium was changed 3 times a week. 2-month-old iNets were dissociated into single-cells suspension using Papain Dissociation System (Worthington, LK003150), passed through a 70-μm cell strainer (Falcon, 07-201-431), resuspended in maturation medium and re-plated (120,000 cells/well) into μ-Plate 96 Well Black (Ibidi 89626).

### Macrophage precursors generation and integration in iNets

Macrophage precursors were generated starting from the KOLF2.1J parental line provided by the iPSC Neurodegenerative Disease Initiative (iNDI) from the NIH’s Center for Alzheimer’s and Related Dementias (CARD) adapting the protocol from van Wilgenburg et al., 2013 (51). In short, iPSCs were seeded into AggreWell 800 plates (STEMCELL Technologies, 34811) pre-treated with Anti-Adherence Rinsing Solution (STEMCELL Technologies, 07010) at a density of 120,000 cells/well to initiate embryoid bodies (EBs) formation. Cells were cultured in StemFlex medium (STEMCELL Technologies, A3349401) supplemented daily with 50 ng/mL BMP4 (Miltenyi Biotec, 130-111-165), 50 ng/mL VEGF (Peprotech, PHC9391), and 20 ng/mL SCF (CellGenix, 1418-050). After four days, EBs were transferred into T75 flasks (Sigma-Aldrich) at approximately 75 EBs per flask and cultured in X-VIVO 15 medium (Lonza, BE02-060F) supplemented with 100 ng/mL M-CSF (Invitrogen, 130-096-492), 25 ng/mL IL-3 (Peprotech, 1402-050), 2 mM GlutaMAX (Gibco 35050-061), 1× Penicillin/Streptomycin (Sigma, P4333-100ML), and 0.055 mM β-mercaptoethanol (Invitrogen, 31350010). Medium was refreshed twice weekly. After 5 weeks of factories set-up, mature macrophage precursors (pMacpre) emerging in the supernatant were collected and cultures replenished with fresh culture media. Harvested pMacpre were then strained through a 40-μm cell strainer (Falcon) and centrifuged at 200g for 5 minutes at room temperature. Cells were resuspended in iNets maturation medium supplemented with 100ng/mL IL-34 (Peprotech) and plated on top of previously subcultured iNets (30,000 cells/well) into μ-Plate 96 Well Black. Two-thirds of the medium was replaced three times per week over 15 days to promote the maturation of pMacpre into microglia-like cells (pMGL).

### Cloning, lentiviral vector production and transduction

The lentiviral transfect vector for the inducible expression of EGFP was generated from the previously described all-in-one monocistronic TetON plasmid (50) by replacing the TDP-43- HA with EGFP sequence via HIFI kit (New Englad Biolabs, E5520S). The lentiviral transfect vector mTRE-EGFP was generated, harvested and concentrated as described previously (50). Lentiviral pellet was resuspended in neural maturation medium containing all supplements except forskolin and cAMP, to obtain 10x concentrated preparations. Lentiviral titer was assessed using Lenti-X GoStix Plus (Takara, 631280). Lentiviral preparations were aliquoted and stored at −80°C until use. For expression of EGFP in iNets, cultures were transduced with 1,000 ng/ml lentivirus in neural maturation medium 18 days after subculturing and 15 days after macrophage precursors addition to the neural network. The day after transduction, complete medium was exchanged to neural maturation medium containing 1 μg/ml doxycycline (Clontech, 631311) to induce transgene expression. Doxycycline was included in subsequent medium refreshments every other day for the following 12 days, when the cultures were fixed for analysis.

### Fixation, immunofluorescence, imaging and quantification of iNets and pMGL

iNets and pMGL were fixed with pre-warmed 16% methanol-free formaldehyde (Pierce, 28908) added directly into the culture medium, diluted to 4% final formaldehyde concentration, and incubated for 20 min at room temperature. Cells were then washed once with PBS (Gibco, 10010015) for 10 min, once with washing buffer (PBS with 0.2% Triton X-100 (Sigma, T9284)) for 10 min and then blocked with 10% normal donkey serum (Sigma-Aldrich, S30-M) and 0.2% Triton X-100 in PBS blocking buffer filtered via stericup (Millipore, S2GPU02RE) for 1 hour at room temperature. Primary antibodies-MAP2 (chicken, Abcam, ab5392, 1:1000), anti-IBA1 (rabbit, Abcam, ab178846, 1:500), anti-GFP (mouse, Proteintech, 66002-1-IG, 1:1000) and anti-human Osteoactivin/GPNMB antibody (Novus Biologicals, AF2550) - were diluted in blocking buffer and incubated overnight at 4 °C on an orbital shaker. The day after, cells were washed 3 × 15 min in washing buffer at room temperature and secondary antibodies - Alexa Fluor 405–conjugated donkey anti-goat IgG (Abcam, ab175664, 1:1000), Alexa Fluor 488– conjugated donkey anti-mouse IgG (Invitrogen, A-21202, 1:1000), Alexa Fluor 568– conjugated donkey anti-rabbit IgG (Invitrogen, A-10042, 1:1000), Alexa Fluor 647–conjugated donkey anti-chicken IgG (Invitrogen, A-78952, 1:1000), were then diluted in blocking buffer and incubated for 1 hour at room temperature. Cells were then again washed 3 × 15 min in washing buffer at room temperature with DAPI (Thermo Scientific 62248) diluted to 1 μg/ml in the final washing buffer wash. Cells were finally washed 1 × 15 min in PBS at room temperature, and PBS was then added to the wells to store the stained cells at 4 °C. Stained cells were imaged using high content scanner MD ImageXpress Confocal HT.ai (40× Water Apo LambdaS LWD objective; 50 μm spinning disk; 2,048 × 2,048 pixels; 30 z-steps per stack and 0.3 μm step size) for quantification. Laser and detector settings were kept the same for each staining and all imaged conditions.

The acquired images were analyzed using custom macros written to suit the FIJI distribution of ImageJ2 (97). To facilitate cell segmentation of pMGLs and iNets, several computations were performed on images capturing IBA1 and MAP2-antibody signals, respectively. First, smoothing via Gaussian blur was performed to reduce noise. Subsequently, background subtraction and the use of an automated intensity threshold enabled the capture of cell candidates in a generated image mask. Moreover, brightness and contrast settings were made uniform for all MAP2 and IBA1 images, respectively. Finally, the segmented regions of interest (ROIs) were filtered using criteria tailored to exclude non-cell objects via filters for size and degree of circularity. Using the resulting mask, average fluorescent intensities were computed per ROI via a second automated script. While only MAP2 signals were used to capture iNets intensities, pMGL intensities were derived from the GPNMB and IBA1-signal channels, both by usage of the IBA1-derived ROIs. This ensured that GPNMB-negative pMGLs were also considered within the image analysis. Furthermore, the GPNMB intensity average per each ROI were divided by the corresponding IBA1 intensity average for normalization, ensuring that technical variation during image acquisition was accounted for.

### Statistical Analysis

For the densitometric analysis of all Western blots, variation in protein expression between NBH and prion conditions across different brain regions was assessed by calculating the standard deviation for each condition, followed by independent two-sample t-tests to evaluate the statistical differences between the NBH and prion-treated groups.

In the analysis of overlapping genes between conditions (Figure 3A and Supplementary Figure 1B), Fisher’s t-test was applied to assess the statistical significance of the overlap between gene sets.

For qPCR analyses, we calculated the standard deviation for each condition to assess gene expression variability. Statistical significance in Figure 3C was evaluated using independent two-sample t-tests to compare *Gpnmb* RNA expression between NBH and prion-inoculated groups across different cell types. In cases where deviations from normality were observed (Figure 2C and Supplementary Figure 9C), such as in the prefrontal cortices of human samples, the Wilcoxon rank-sum test was used to compare fold changes between sCJD and control patients.

In Figure 6C and Supplementary Figure 15B, pairwise comparisons between experimental conditions were performed using two-sided Wilcoxon rank-sum tests. P-values were adjusted for multiple testing using the Benjamini-Hochberg method.

For phagocytosis assay analysis (Figure 6E), we calculated the standard deviation for each condition, followed by ANOVA and Dunnett’s post-hoc correction.

For differential expression analysis, we employed DESeq2, a tool commonly used for RNA-seq data analysis. DESeq2 was used to compare gene expression levels between conditions, with p-values adjusted by the Benjamini-Hochberg method to control the false discovery rate (FDR). Genes with adjusted p-values below 0.05 were considered significantly differentially expressed and log2 fold change values quantified the magnitude of expression changes.

For over-representation analysis (Figure 3B and Supplementary Figure 2), a custom function was used to identify enriched biological processes associated with differentially expressed genes. Fisher’s exact test was employed to determine whether specific GO terms were over-represented in the gene list and p-values were adjusted using the Benjamini-Hochberg method to control FDR.

In Supplementary Figure 10A, KEGG pathway enrichment analysis was conducted for upregulated genes using a p-value cutoff of 0.05. Fisher’s exact test was applied to identify over-represented pathways.

For mice weight analysis (Supplementary Figure 9C), mean weights for each condition across weeks were calculated to track weight trends over time. Independent two-sample t-tests were performed weekly to determine statistical differences in weight between the prion and control groups.

For ELISA-related analyses (Figure 2D-E), protein levels between conditions were compared by calculating the standard deviation of OD values, with statistical significance assessed via the Wilcoxon rank-sum test for each organ.

## Data Availability

Raw sequencing data as well as processed data from this manuscript are freely available via GEO accession number GSE277577. Code to reproduce results generated in this manuscript are available at https://github.com/dcare91/ST_prions. Raw immunofluorescent microscopy images are publicly available via https://doi.org/10.5281/zenodo.15824298.

## Author’s Contributions

DC conceptualized the study, performed bioinformatic analyses (including preprocessing, alignment, and visualization of spatial transcriptomics data), and conducted the experiments. He secured funding, created all figures and tables, and was responsible for drafting the initial manuscript, integrating co-author feedback, and ensuring the overall coherence of the study. GM performed Western blot analysis for Spp1, Lgals3, Vim, and NeuN in various brain regions of NBH and RML6 mice and conducted BV2-Gpnmb knockout experiments to assess phagocytosis activity. He also carried out BV2 cell treatments with conditioned media from prion-infected CAD5 cells and contributed to writing the original manuscript. Additionally, he prepared brain slices for spatial transcriptomics and performed the 10x library preparation for all samples. LP conducted immunofluorescence staining of brain slices for Gpnmb and Lgals3 co-localization and contributed to BV2 cell experiments with apoptotic-conditioned media. She also optimized the FACS protocol for apoptotic body isolation and conditioned media collection. She performed the imaging analysis of human neuron-microglia coculture and also contributed significantly to writing the materials and methods section. MC performed immunohistochemistry experiments, assisted in establishing the spatial transcriptomics protocol, and played a key role in developing CNS cell-type sorting and Gpnmb qPCR protocols. She also contributed significantly to writing the materials and methods section. BG performed the validation of apoptosis-induced Gpnmb upregulation in the neuro-immune co-culture system and carried out the subsequent staining and imaging. YL maintained cell lines and conducted Western blot analyses for Gpnmb, Spp1, Lgals3, and Vim in cell-based experiments. ME contributed to the development and optimization of Gpnmb ELISA protocols and conducted the related data analysis. MHP and MP supervised BG in the experimental setup, providing structured guidance, reagents, and technical suggestions. MM and IZ collected, diagnosed, and provided CJD patient samples, which were used by ME for ELISA-based GPNMB fragment measurements. ED provided critical feedback on project design and conducted early experiments that were not included in the final manuscript. She was responsible for drafting the initial manuscript, integrating co-author feedback, and ensuring final manuscript coherence. AA conceptualized the study, provided animals, personnel, and IT resources, oversaw the research team, offered experimental guidance, managed the project, secured funding, contributed to manuscript drafting, and thoroughly reviewed the final version.

## Acknowledgements

We thank Petra Schwarz and Laura Cervantes for their work in maintaining the animals and Maruša Koderman, Nicola Conneely, Lidia Madrigal and Marija Mitrovic for assistance. Andrea Armani provided invaluable support in ensuring the manuscript’s coherence and logical flow. We are grateful to the Functional Genomic Center of Zurich (FGCZ) for performing the spatial transcriptomics sequencing. Special thanks go to Regina Reimann and Irina Abakumova for their discussions and advice regarding human patient data. We also extend our gratitude to Giuseppe Legname’s group, particularly Lea Nikolic, Marco Zattoni and Marika Mearelli, for providing us with cDNA samples from sCJD patients and sex- and age-matched controls.

## Financial Disclosure

A.A. is supported by institutional core funding by the University of Zurich, a Distinguished Scientist Award of the NOMIS Foundation and grants from the GELU Foundation, the Swiss National Science Foundation (SNSF grant ID 179040 and grant ID 207872, Sinergia grant ID 183563), the Human Frontiers Science Program (grant ID RGP0001/2022), the Michael J. Fox Foundation (grant ID MJFF-022156) and the CJD Foundation. D.C. is supported by a CANDOC grant of the University of Zurich.

---

**Supplementary Figure 1.**
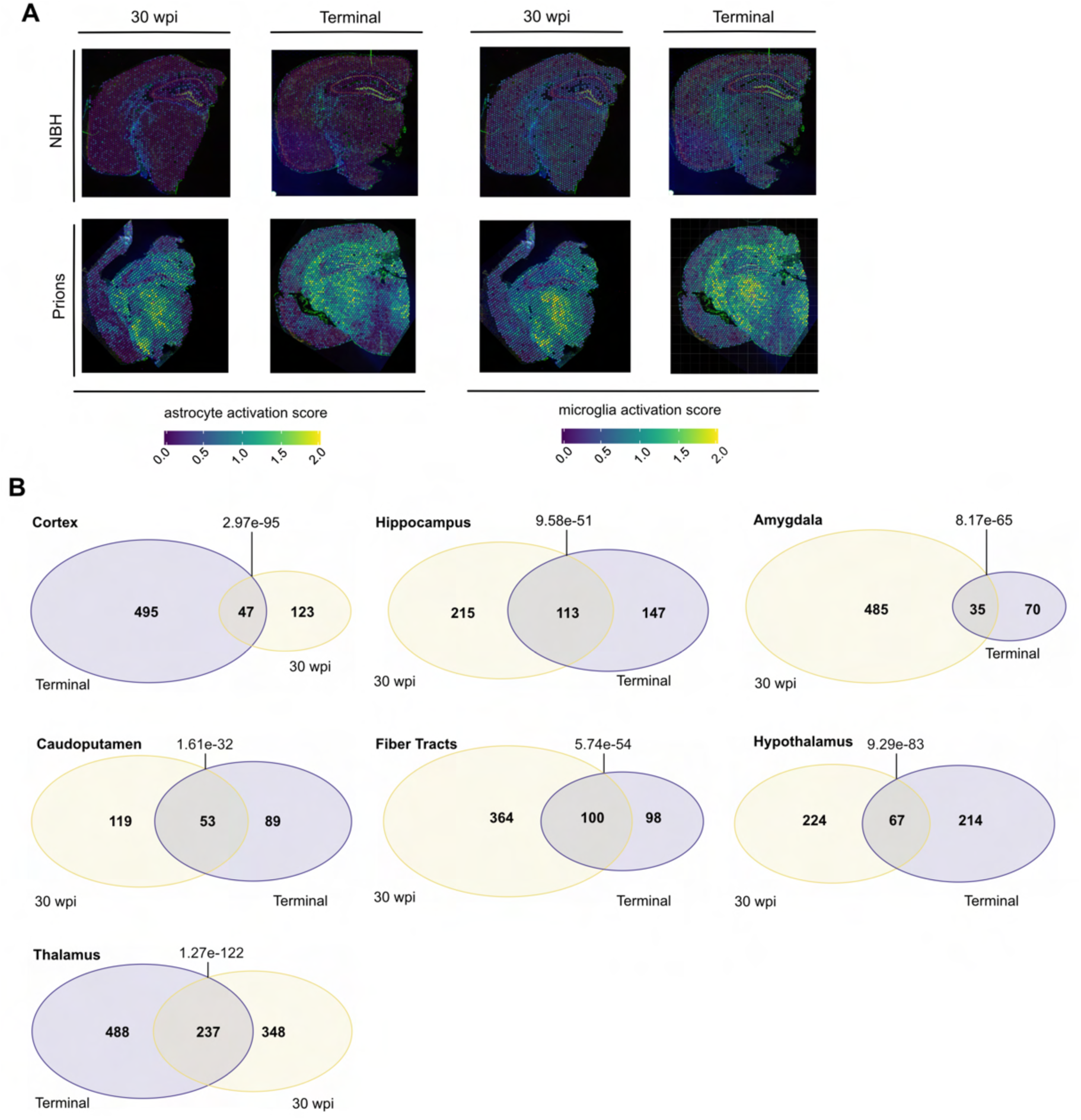
**A)** Panels display results from a 10x Genomics spatial transcriptomic assay, comparing brain slices under two conditions (NBH and Prions) at two time points (30 weeks post-infection (wpi) and Terminal). These images are color-coded to illustrate levels of astrocyte and microglia activation, with activation scores shown at the bottom of the panel. The activation levels are indicated by the color spectrum, where cooler colors (blues) represent lower activation and warmer colors (yellows) indicate higher activation. **B)** This figure illustrates the overlap of differentially expressed genes (DEGs) between two critical disease stages - 30 weeks post-inoculation (wpi) and the terminal stage - across various brain regions including the cortex, hippocampus, amygdala, caudoputamen, fiber tracts, thalamus and hypothalamus. Each Venn diagram represents a specific brain region, detailing the number of unique and shared DEGs at each stage.

**Supplementary Figure 2.**
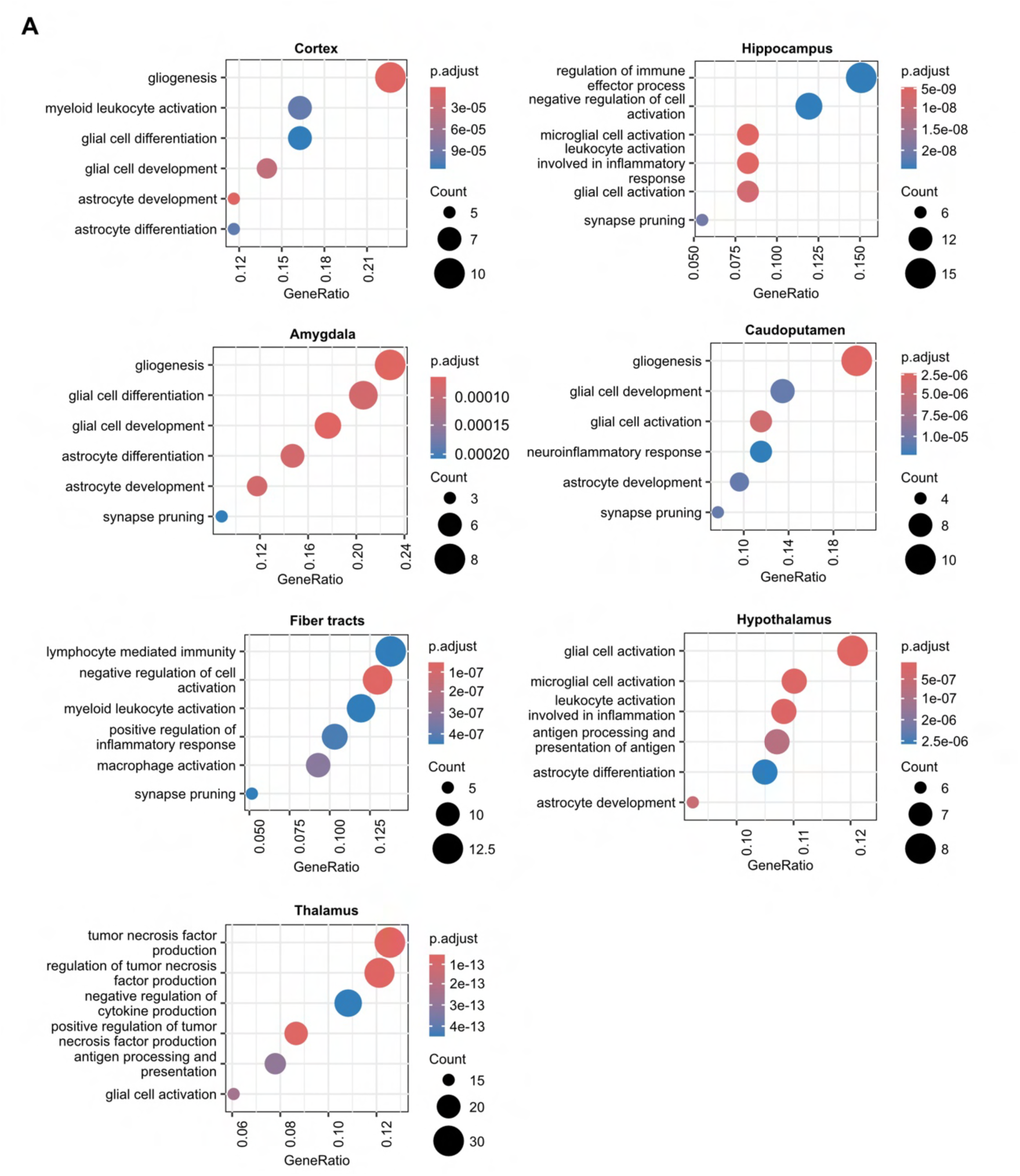
This figure presents an over-representation analysis of genes that overlap between the 30 weeks post-inoculation and terminal stages of PrD across various brain regions. Each panel represents a different region, including the cortex, hippocampus, amygdala, caudoputamen, fiber tracts, hypothalamus and thalamus. Colored circles indicate the enrichment of specific biological processes such as gliogenesis, immune response and synaptic pruning, with colors reflecting p-adjust values and circle size indicating the gene count involved in each process.

**Supplementary Figure 3.**
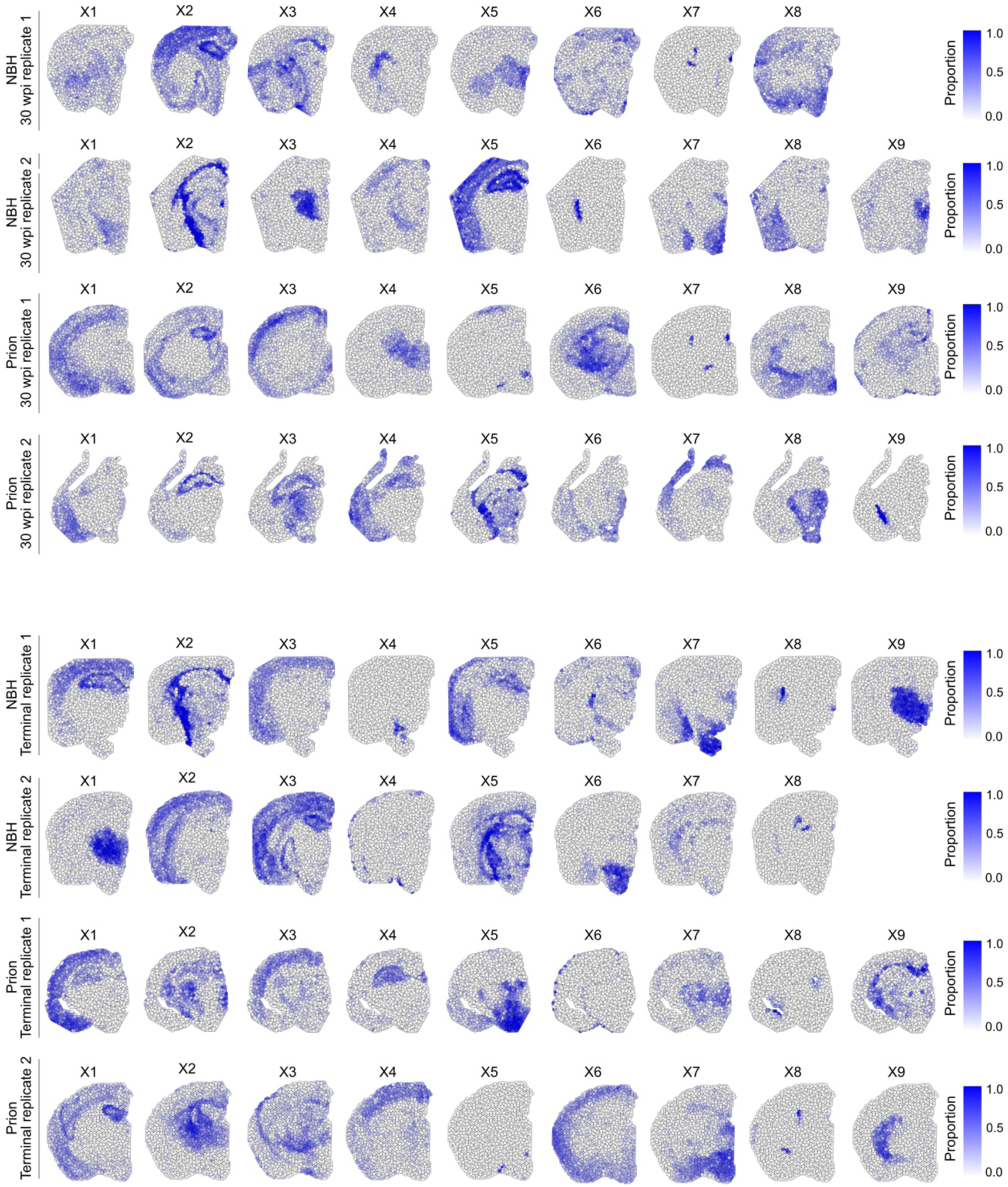
This figure displays the spatial distribution and relative abundance of various cell types identified through gene expression profiles in brain sections from both control (NBH) and prion-infected samples at 30 wpi and the terminal stages. Each row corresponds to a different experimental condition and replicate, mapping the distribution of identified cell types from X1 to X9. The intensity of the color within each brain section indicates the proportion of specific cell types, with darker shades denoting a higher prevalence.

**Supplementary Figure 4.**
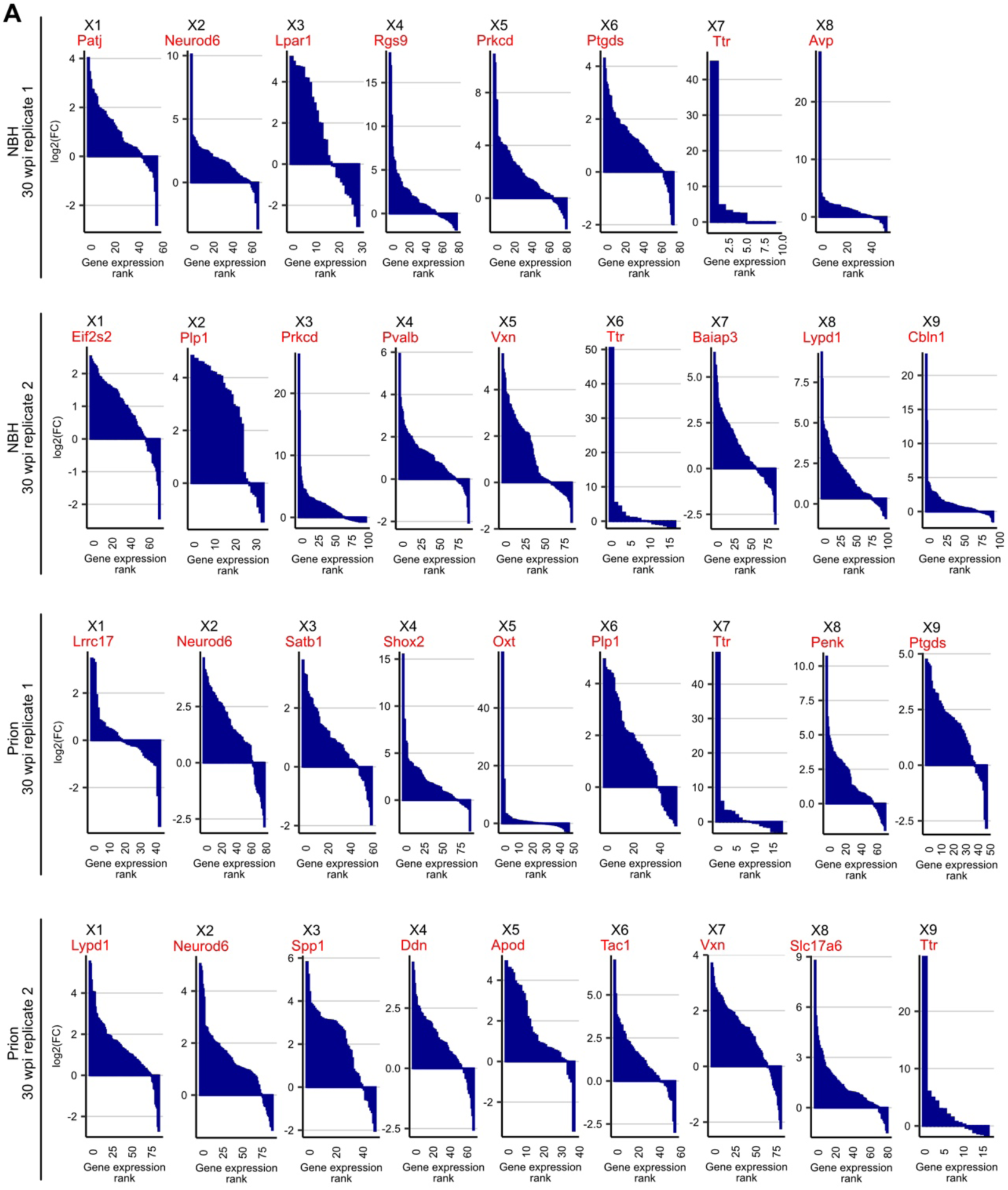
This figure displays the ranked gene expression profiles within identified cell types from both control (NBH) and prion-infected samples at 30 wpi. Each plot, labeled X1 through X8 or X9, corresponds to a specific cell type, showcasing the distribution of gene expression rankings. The plots are named after the most enriched gene (in red) within each identified cell type, which highlights the defining characteristic of that cell type according to the STdeconvolve algorithm. The x-axis of each plot shows the rank of genes within that cell type, sorted by expression level, while the y-axis depicts the log2 fold change of these genes. Different rows reflect distinct experimental conditions and replicates.

**Supplementary Figure 5.**
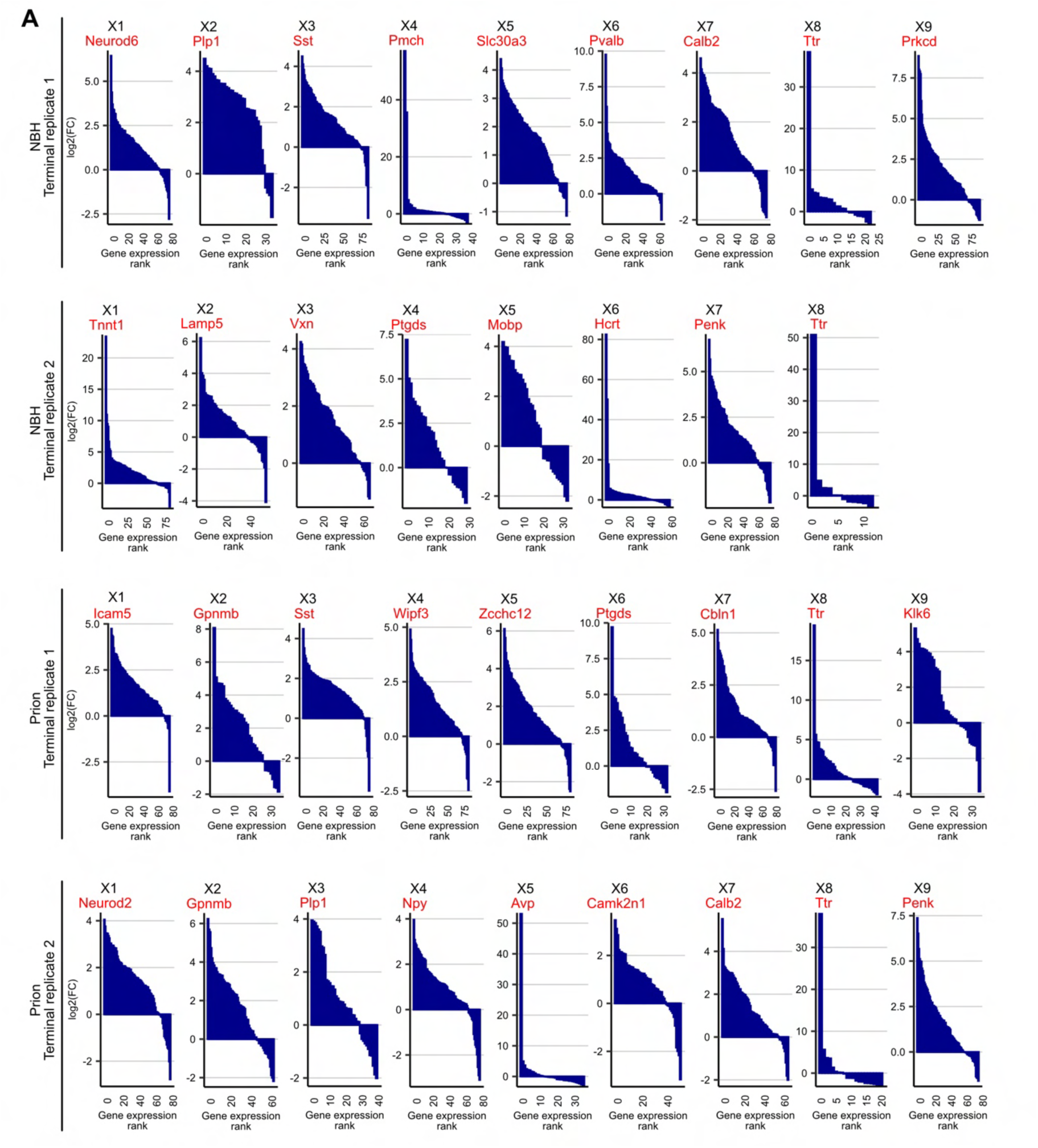
This figure displays the ranked gene expression profiles within identified cell types from both control (NBH) and prion-infected samples at terminal stage of the disease. Each plot, labeled X1 through X8 or X9, corresponds to a specific cell type, showcasing the distribution of gene expression rankings. The plots are named after the most enriched gene (in red) within each identified cell type, which highlights the defining characteristic of that cell type according to the STdeconvolve algorithm. The x-axis of each plot shows the rank of genes within that cell type, sorted by expression level, while the y-axis depicts the log2 fold change of these genes. Different rows reflect distinct experimental conditions and replicates.

**Supplementary Figure 6.**
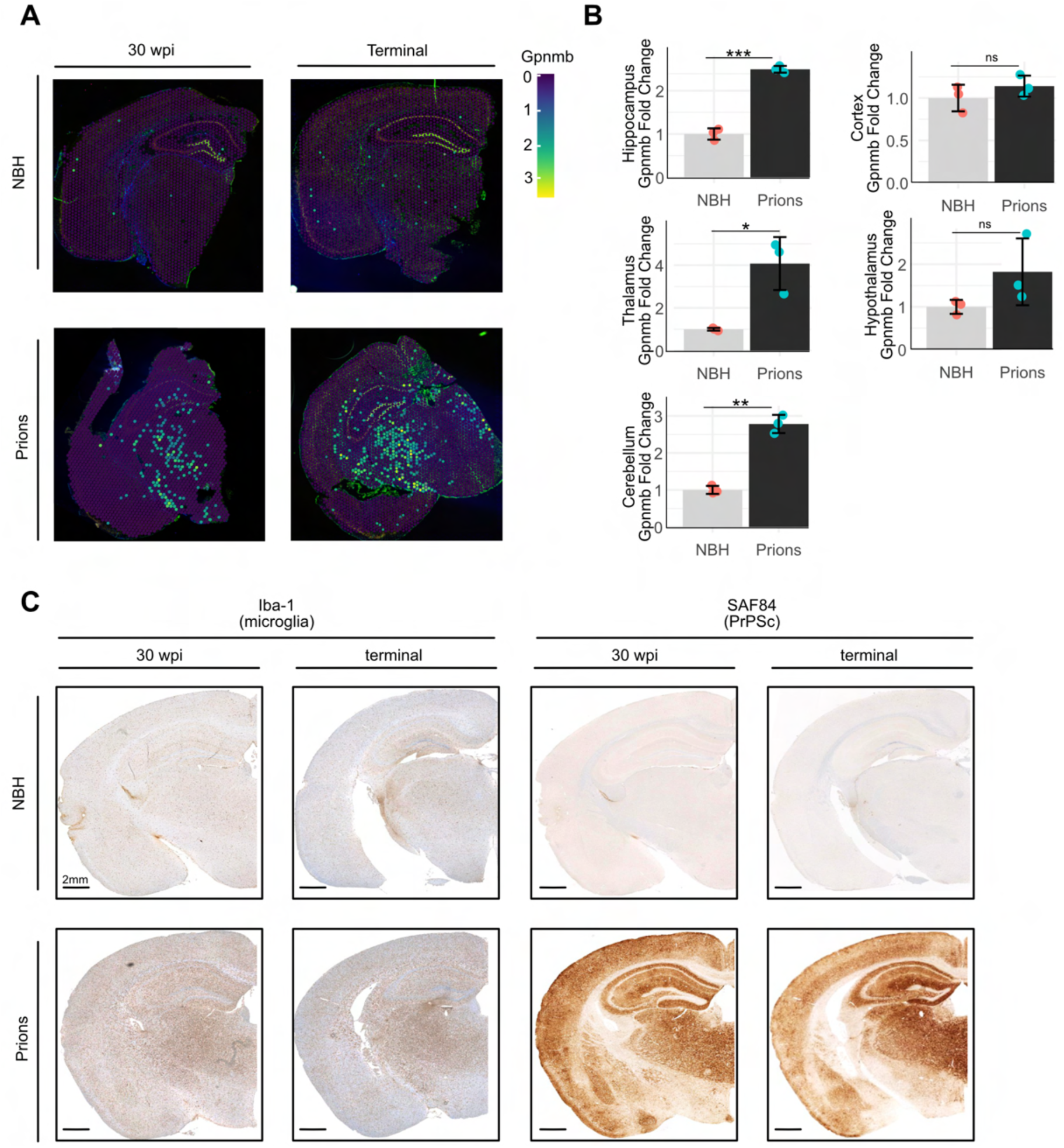
**A)** Panels display results from a spatial transcriptomic assay, comparing brain slices under two conditions (NBH and prions) at two time points (30 wpi and terminal) of a further biological replicate. These images are color-coded to illustrate levels of *Gpnmb* expression, with expression level indicated by the color spectrum, where cooler colors (blues) represent lower activation and warmer colors (yellows) indicate higher activation. **B)** Densitometry (arbitrary densitometry unit, ADU) quantification of the Western Blot in Figure 2B. Statistical significance (*p < 0.05, **p < 0.01, ***p < 0.005, ****p < 0.001) is indicated by asterisks. **C)** Immunohistochemistry of brain sections stained for Iba-1 (a marker for microglia) and SAF84 (a marker for PrP^Sc^). Representative images from control (NBH) and prion-infected mice at 30 wpi) and terminal stages are shown. The sections stained with Iba-1 reveal the presence and activation status of microglia, while SAF84 staining highlights the accumulation of prion protein (PrP^Sc^) in the infected brain tissues. The scale bar represents 2 mm.

**Supplementary Figure 7.**
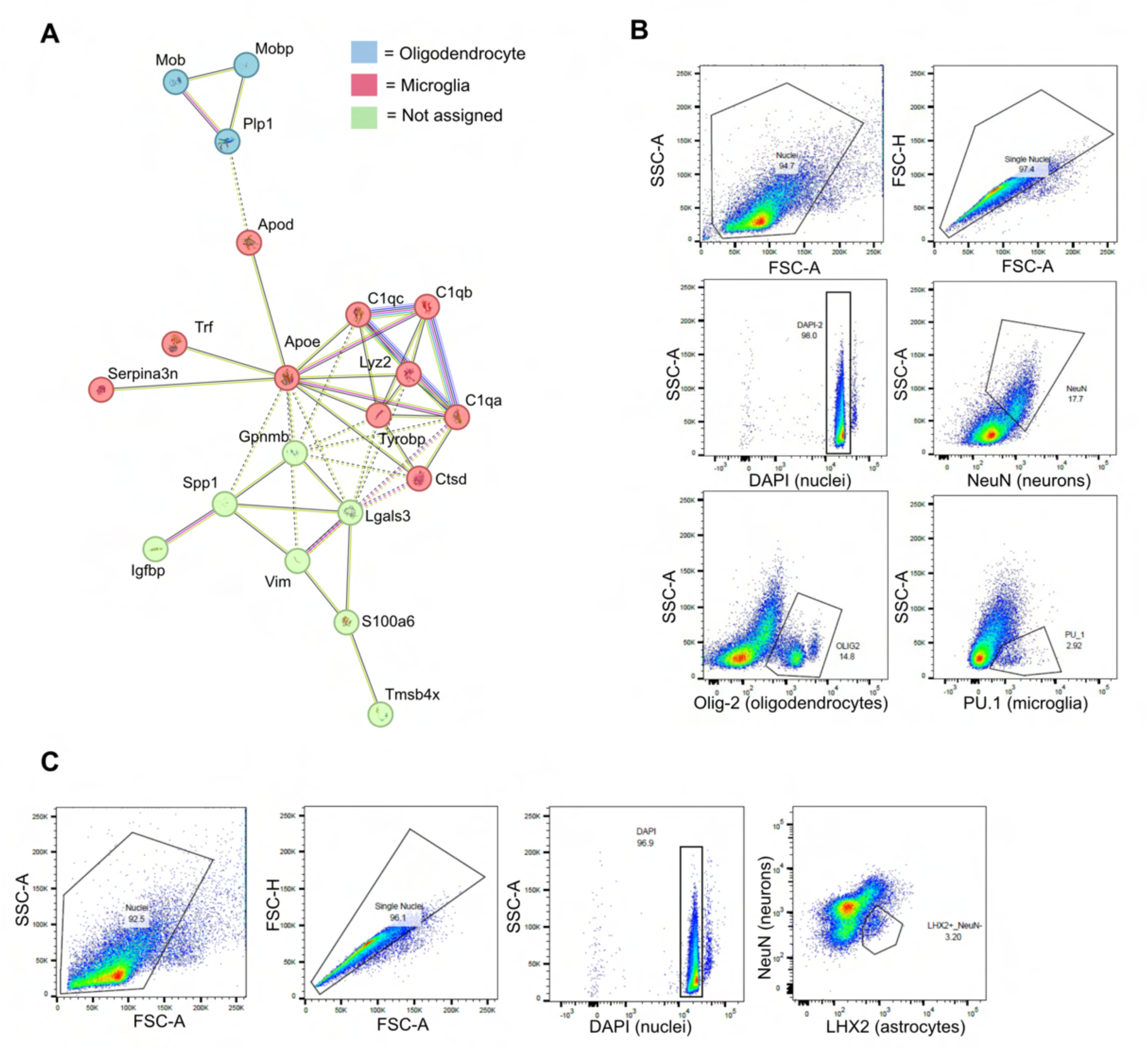
**A)** STRING network of genes associated with the Gpnmb* microglial cell type. Nodes represent individual genes, and edges (lines) depict interactions between these genes. The nodes are color-coded to represent different gene categories: oligodendrocyte-assigned (light blue), microglial-assigned (red), and unassigned (green). **B-C)** These panels display flow cytometry plots for the isolation and characterization of nuclei from prion-infected and control (NBH) brain samples. B) The first plot shows the selection of nuclei based on forward and side scatter (FSC-A vs. SSC-A). The second plot refines the selection to single nuclei (FSC-H vs. FSC-A). The third plot confirms DAPI-positive nuclei (Vio 450 vs. SSC-A). Subsequent plots depict the gating for NeuN^+^ (neuronal nuclei), OLIG2^+^ (oligodendrocyte nuclei), and PU.1^+^ (microglial nuclei). C) Similar gating strategy as for A: selection of nuclei, single nuclei, DAPI-positive nuclei. The final plot shows the gating for NeuN- and LH2^+^ (astrocyte nuclei).

**Supplementary Figure 8.**
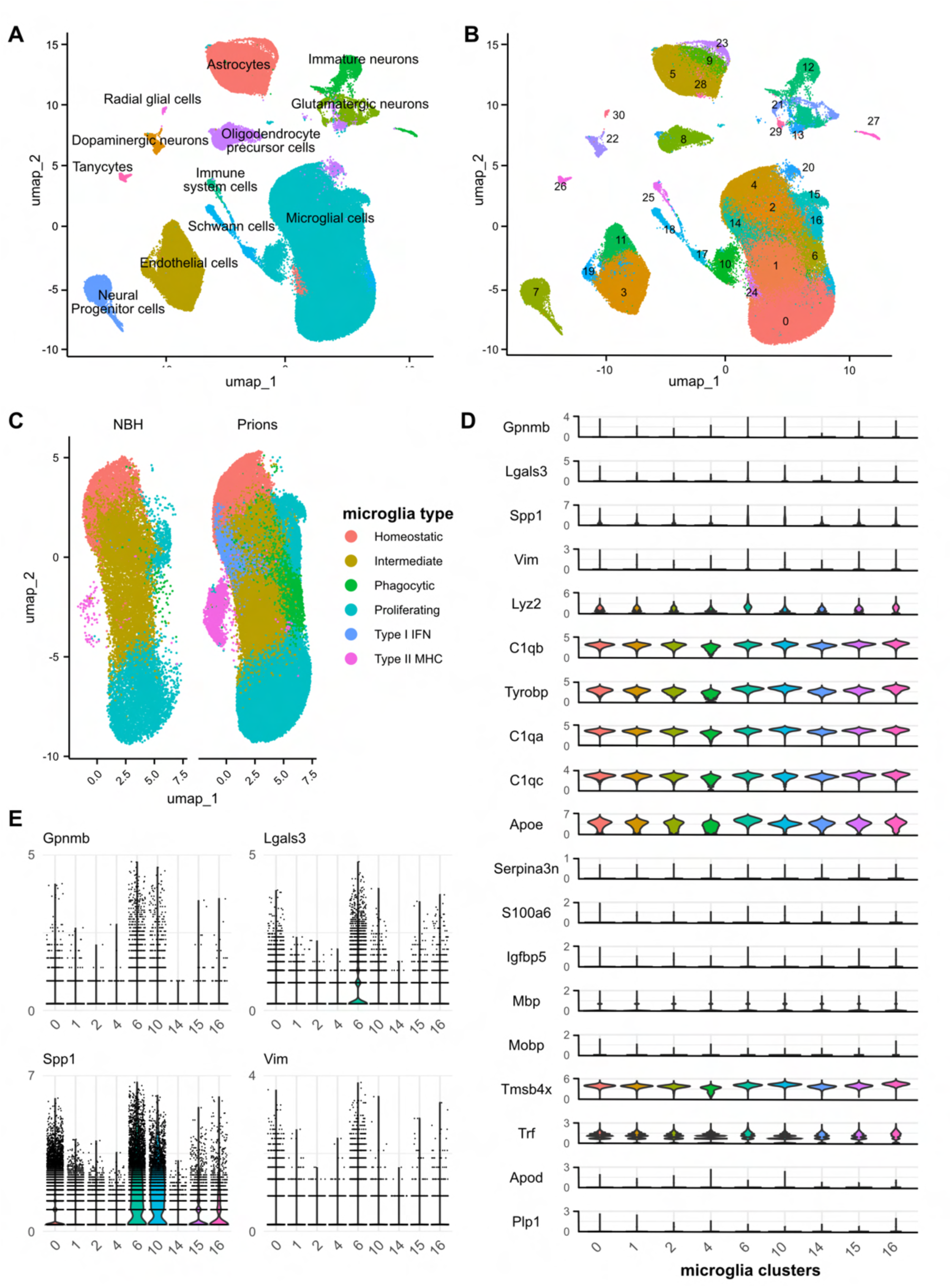
This figure presents a reanalysis of single-cell RNA sequencing data of terminally ill prion infected mice from Slota et al., 2022. **A)** UMAP plots illustrating the distribution of major CNS cell types, including microglia, astrocytes, neurons, oligodendrocyte precursor cells and others. **B)** UMAP plots showing clustering of major CNS cell types into distinct subtypes, with labels indicating specific clusters. **C)** Comparative UMAP plots of microglial cells from NBH and prion-infected conditions, categorized into homeostatic, intermediate, phagocytic, proliferating, type I IFN and type II MHC microglia. **D)** Violin plots displaying the expression levels of the 19 genes obtained from the overlap of the two prion-related replicates by using STdeconvolve statistical model. **E)** Dot plots showing gene expression levels for *Gpnmb*, *Lgals3*, *Spp1* and *Vim* across microglial clusters.

**Supplementary Figure 9.**
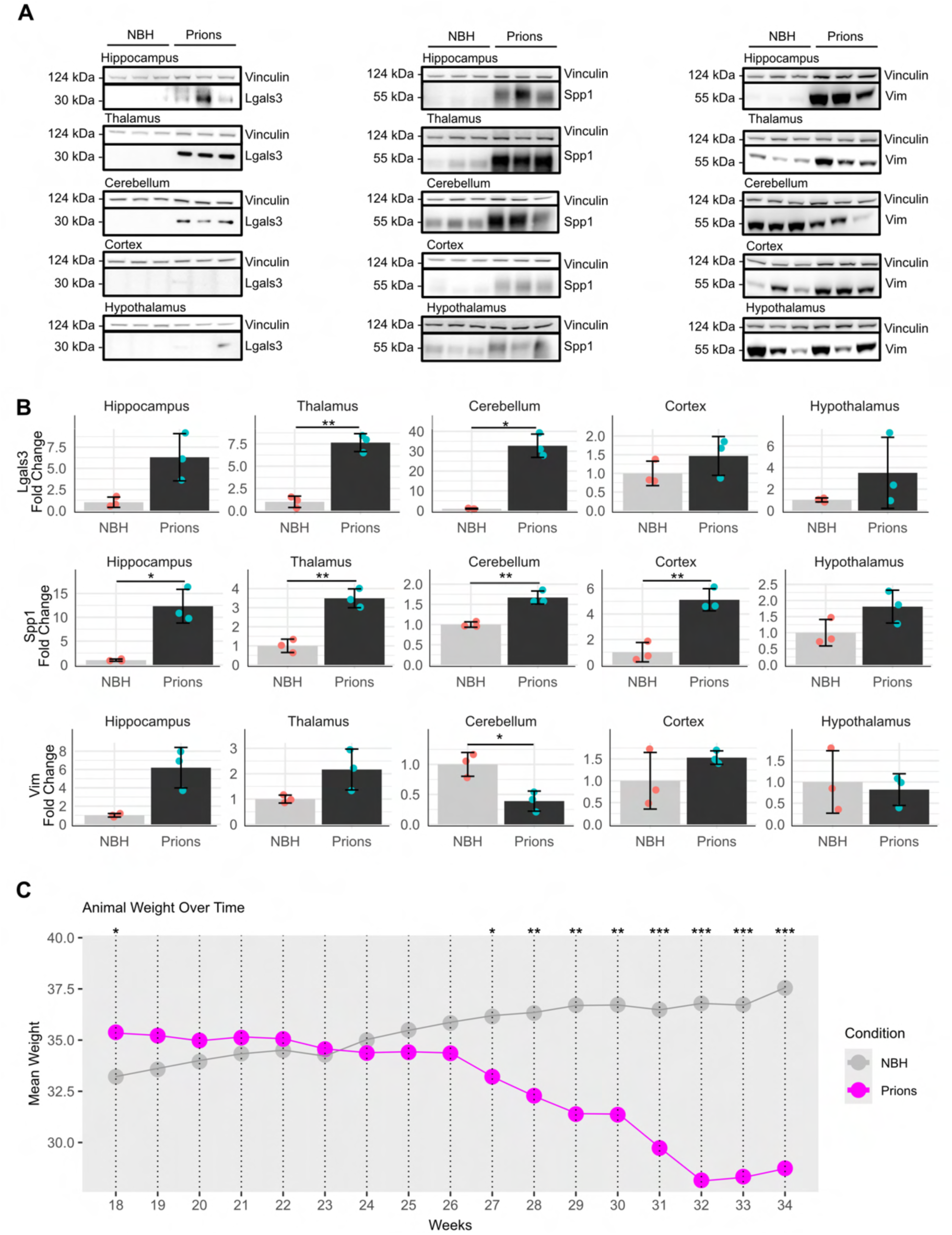
**A)** Western blot analysis showing the protein levels of Lgals3, Spp1, and Vim across different brain regions in control (NBH) and prion-infected samples assessed at the terminal stage. Vinculin is used as a loading control. **B)** Densitometry analysis of the Western blots in panel A, quantifying fold changes in protein expression levels between NBH and prion-infected brains. Significant differences are indicated by asterisks (* p<0.05, ** p<0.01, *** p<0.001), with data presented as mean ± SEM. **C)** Linechart of the mean of (10 biological replicates) mouse weights (y-axis) intraperitoneally treated with NBH (grey) or infected with prions (magenta) assessed by weeks post inoculation (x-axis). Significant differences are indicated by asterisks (* p<0.05, ** p<0.01, *** p<0.001).

**Supplementary Figure 10.**
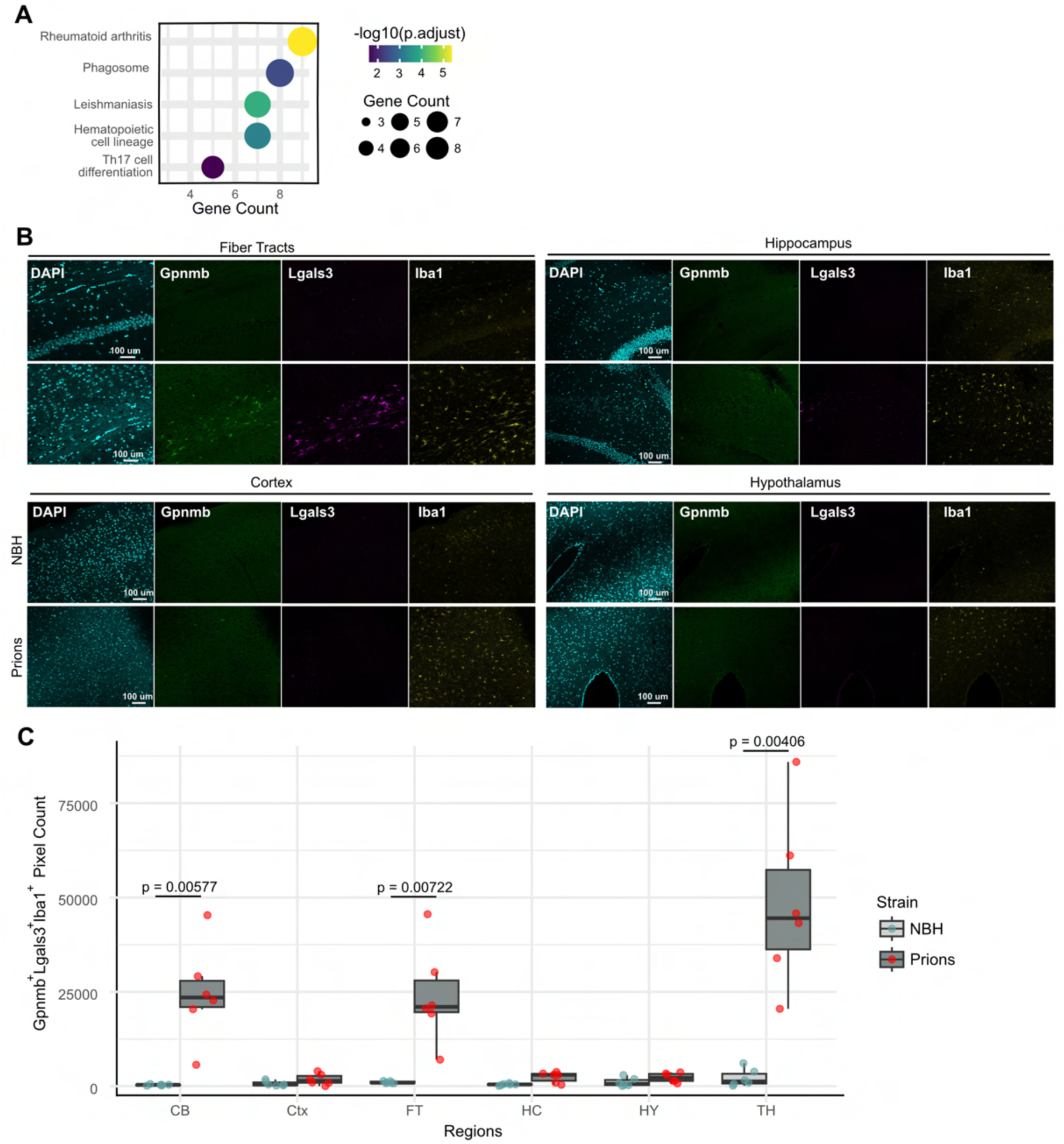
**A)** KEGG enrichment of upregulated genes in Gpnmb⁺/Phagocytic Cluster 6⁺ versus Gpnmb⁻/Phagocytic Cluster 6⁺ microglia from scRNA-seq data (Slota et al. 2022). Dot size indicates gene count, color reflects p-value (red = most significant) **B)** Immunofluorescence targeting DAPI (cyan), Gpnmb (green), Lgals3 (magenta) and Iba1 (yellow) on terminally ill prion infected mice. Two ROI of different brain regions are shown. **C)** Boxplot comparing the positive pixel count across different conditions. Each box represents the distribution of positive pixels for a given condition, with custom colors distinguishing the groups. Dots for each regional condition (CB = cerebellum, Ctx = cortex, FT = fiber tracts, HC = hippocampus, HY = hypothalamus, TH = thalamus) represent two technical replicates from three biological replicates (n = 3). The central line in each box indicates the median, while the upper and lower bounds represent the interquartile range (IQR). Whiskers extend to 1.5 times the IQR and outliers are shown as individual points. Significant differences between groups are indicated by asterisks (* p<0.05, ** p<0.01, *** p<0.001).

**Supplementary Figure 11.**
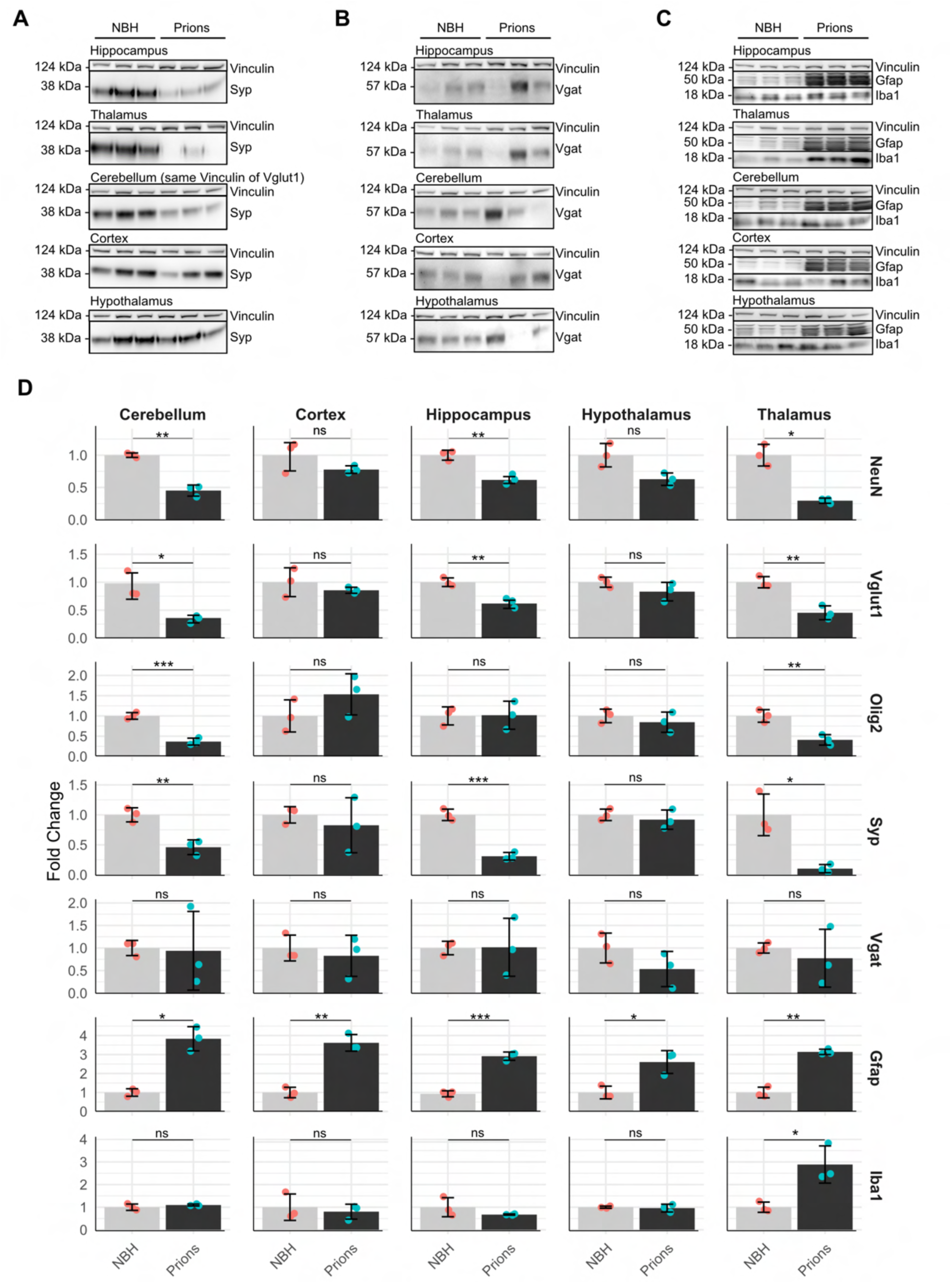
Western blot analysis showing the protein levels of **A)** Syp, **B)** Vgat, **C)** Gfap and Iba1 across different brain regions in control (NBH) and prion-infected samples assessed at the terminal stage. Vinculin is used as a loading control. **D)** Densitometry analysis of the Western blots in Figure 5D and Supplementary Figure 11A-C, quantifying fold changes in protein expression levels between NBH and prion-infected brains. Significant differences are indicated by asterisks (* p<0.05, ** p<0.01, *** p<0.001), with data presented as mean ± SEM.

**Supplementary Figure 12.**
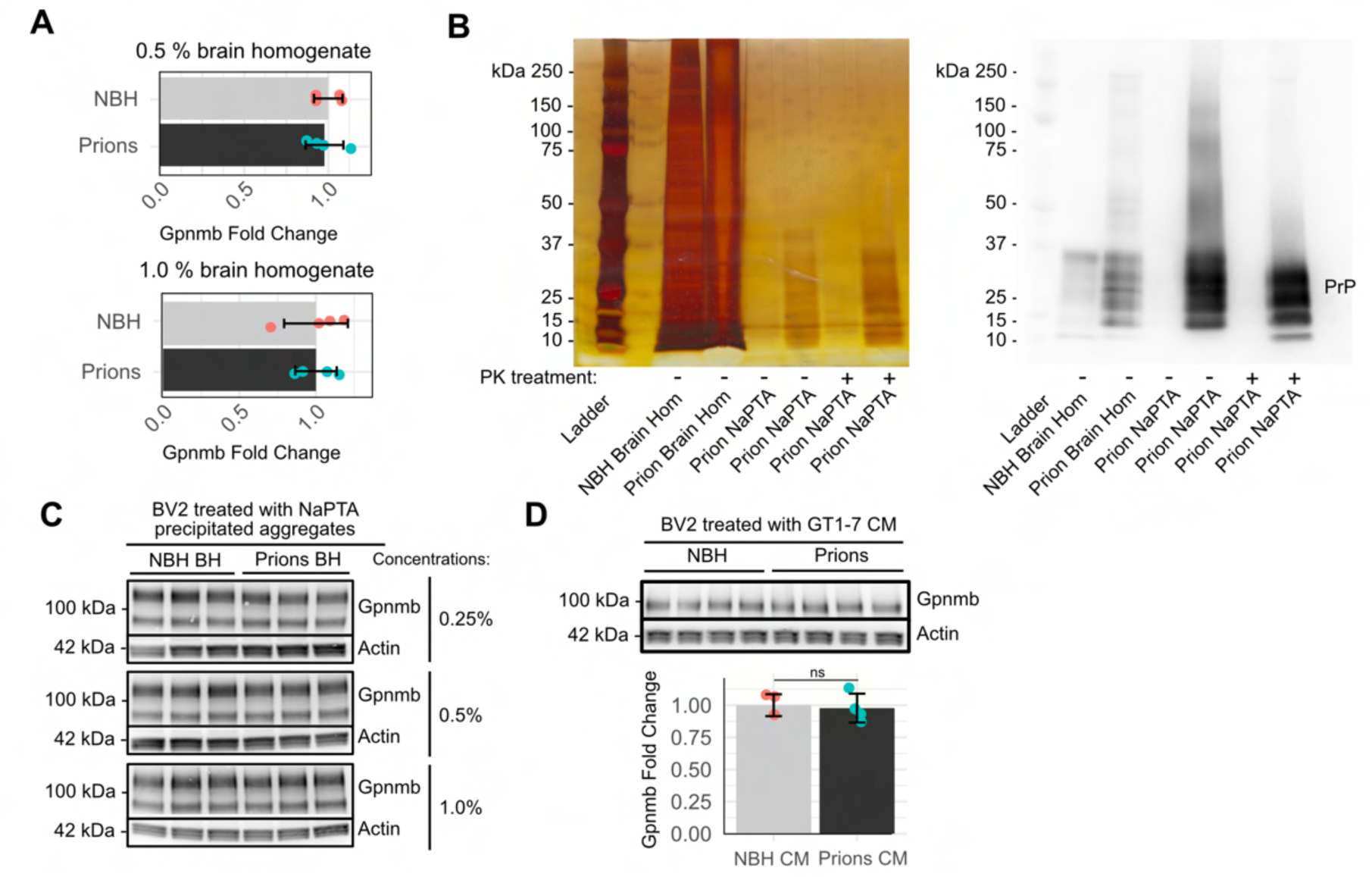
**A)** qPCR of Gpnmb expression in BV2 cells treated with 0.5% and 1.0% brain homogenates from NBH and RML6 prion-infected mice. **B)** On the left, uncut silver-stained 16% SDS-PAGE gel displays both normal brain homogenate (NBH) and brain homogenate from prion-infected mice alongside purified fractions with NaPTA treatment. On the right, uncut western blot displays the same condition stained with anti-PrP antibody (POM1). The “+” and “–” symbols indicate the presence or absence of proteinase K (PK) treatment during purification. **C)** Gpnmb- and Actin-stained western blots of BV2 treated with NaPTA isolated aggregates from both NBH and prion-containing brain homogenates are shown. The equivalent of 0.25%, 0.5%, and 1% (w/v) brain homogenate from both NBH and prion-infected samples was used to load NaPTA-precipitated aggregates. **D)** Western blot and densitometric analysis of Gpnmb in BV2 cells treated for 48 hours with conditioned media (CM) from prion-infected GT1-7 cells. Significant differences between groups are indicated by asterisks (* p<0.05, ** p<0.01, *** p<0.001).

**Supplementary Figure 13.**
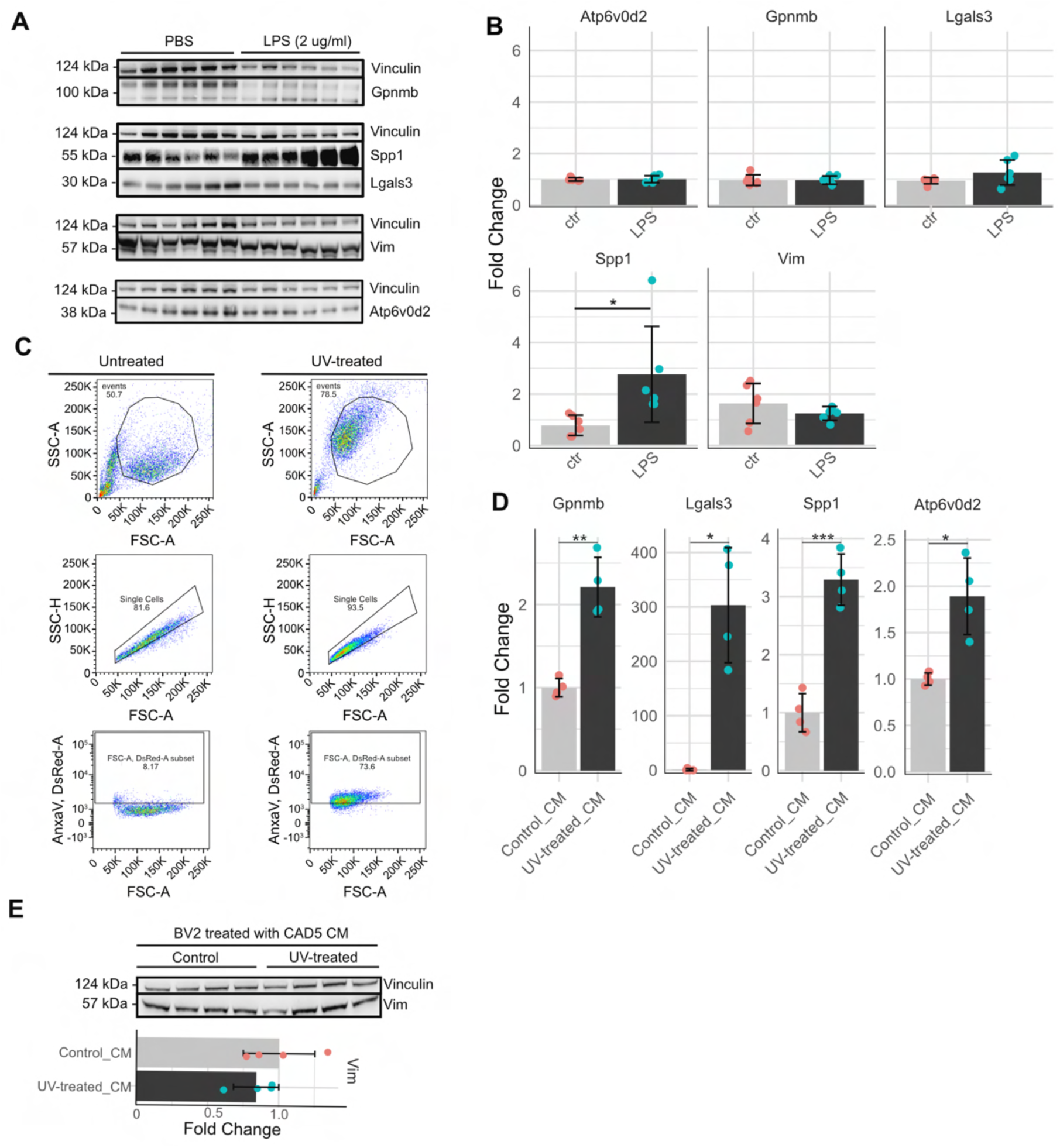
**A)** Western blot of BV2 cells treated with PBS or LPS (2 μg/ml) for 24 hours, probing Gpnmb, Spp1, Lgals3, Vim, Atp6v0d2 and Vinculin. **B)** Densitometric analysis of panel E, showing fold change in protein expression (PBS vs. LPS). **C)** FACS analysis of Annexin-V (AnxaV) in UV-treated vs. untreated CAD5 cells confirms apoptosis. **D)** Densitometric analysis of Western blot (Figure 6A) in BV2 cells cultured with CM from UV-treated and untreated CAD5 cells. **E)** Top: Western blot of BV2 cells treated with CM from UV-treated CAD5 cells, probing Vim vs. Vinculin. Bottom: Densitometric analysis. Significant differences between groups are indicated by asterisks (* p<0.05, ** p<0.01, *** p<0.001).

**Supplementary Figure 14.**
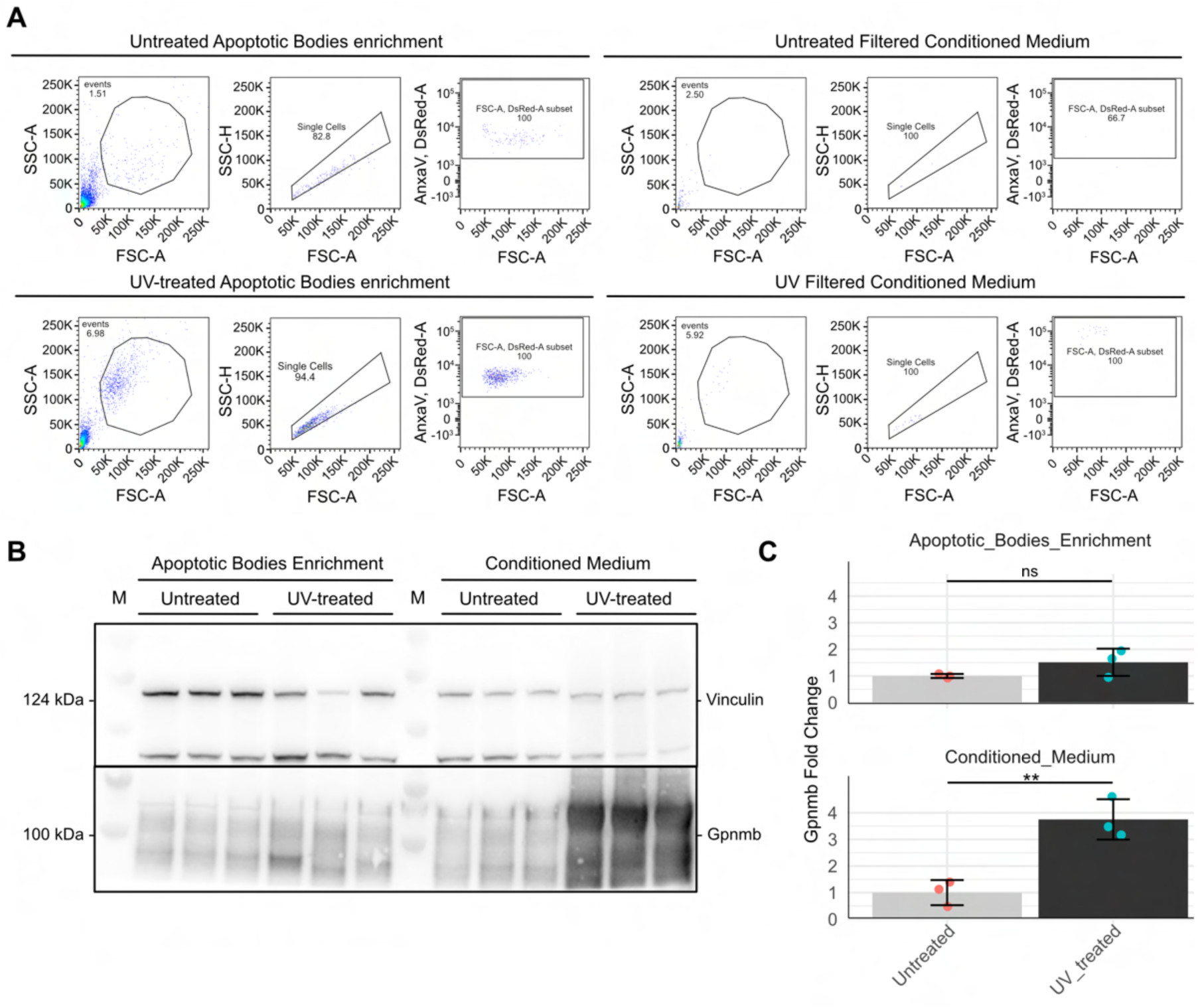
**A)** Flow cytometry analysis of apoptotic bodies and filtered conditioned medium from untreated and UV-treated CAD5 cell line. Representative forward scatter (FSC-A) versus side scatter (SSC-A) plots show particle size distribution. Gating strategies identify single cells and annexin V-positive apoptotic bodies. **B)** Uncut western blot of Gpnmb protein levels in BV2 cell line treated with apoptotic bodies and conditioned medium from CAD5 cell line untreated and UV-treated samples. Vinculin serves as a loading control. **C)** Quantification of Gpnmb protein levels from panel B, shown as fold change relative to untreated samples. Significant differences between groups are indicated by asterisks (* p<0.05, ** p<0.01, *** p<0.001).

**Supplementary Figure 15.**
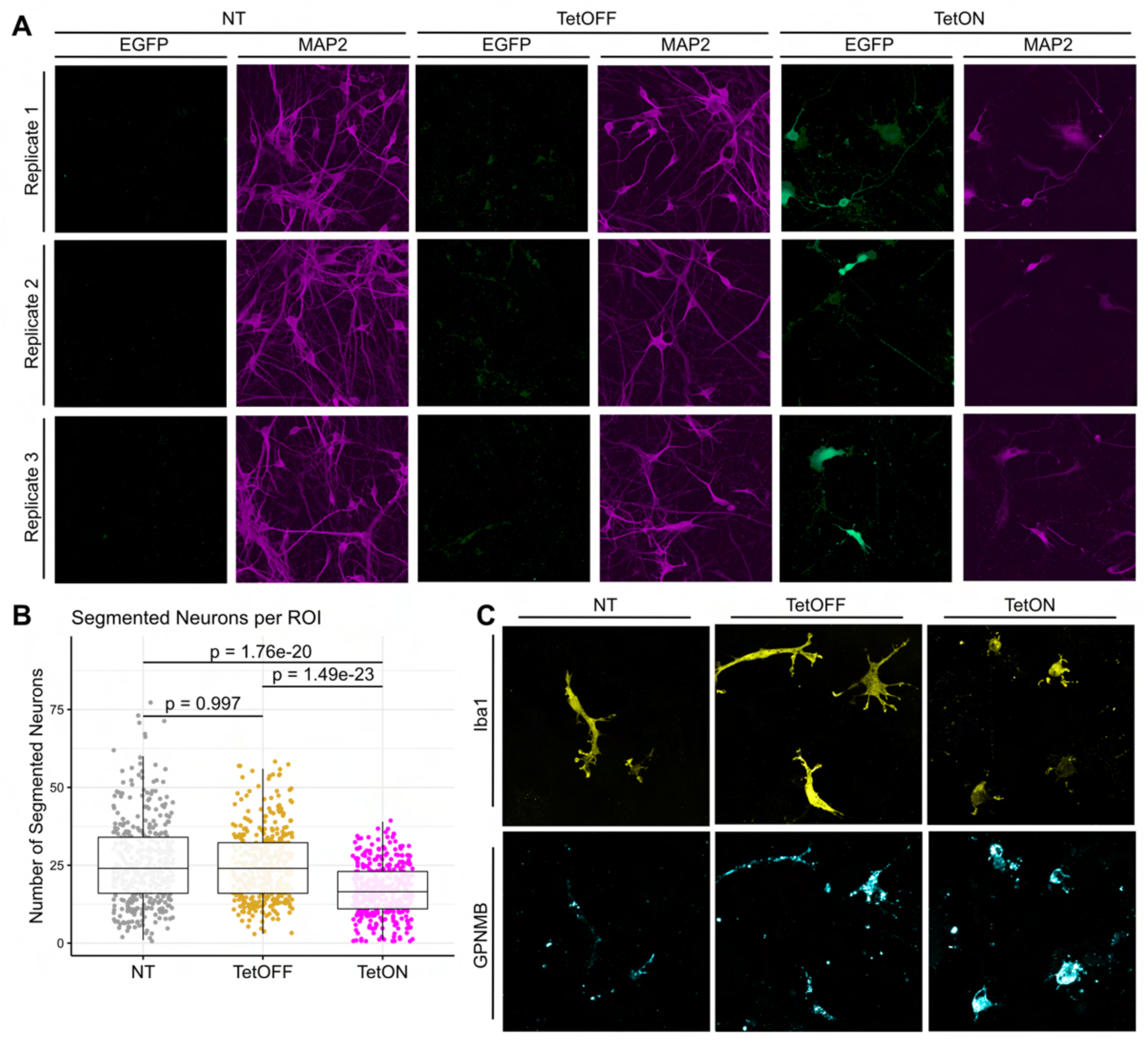
**A)** Representative immunofluorescence images acquired at 40x magnification from three biological replicates of human co-cultures comprising iNets and microglia. EGFP and MAP2 channels are shown separately to visualize neuronal structure and health. NT (non-transduced control), TetOFF, and TetON conditions are displayed side-by-side. All images were acquired under identical settings and processed equally for display. **B)** Boxplots show the number of segmented neurons per region of interest (ROI) in the NT (non-treated), TetOFF (doxycycline withdrawn), and TetON (doxycycline administered) conditions (data represented in Figure 6B). Points represent individual MAP2-derived ROIs pooled from two biological experiments. **C)** Representative confocal images acquired at 63x magnification showing Iba1 (yellow) and GPNMB (cyan) immunostaining in the NT, TetOFF, and TetON conditions.

**Supplementary Figure 16.**
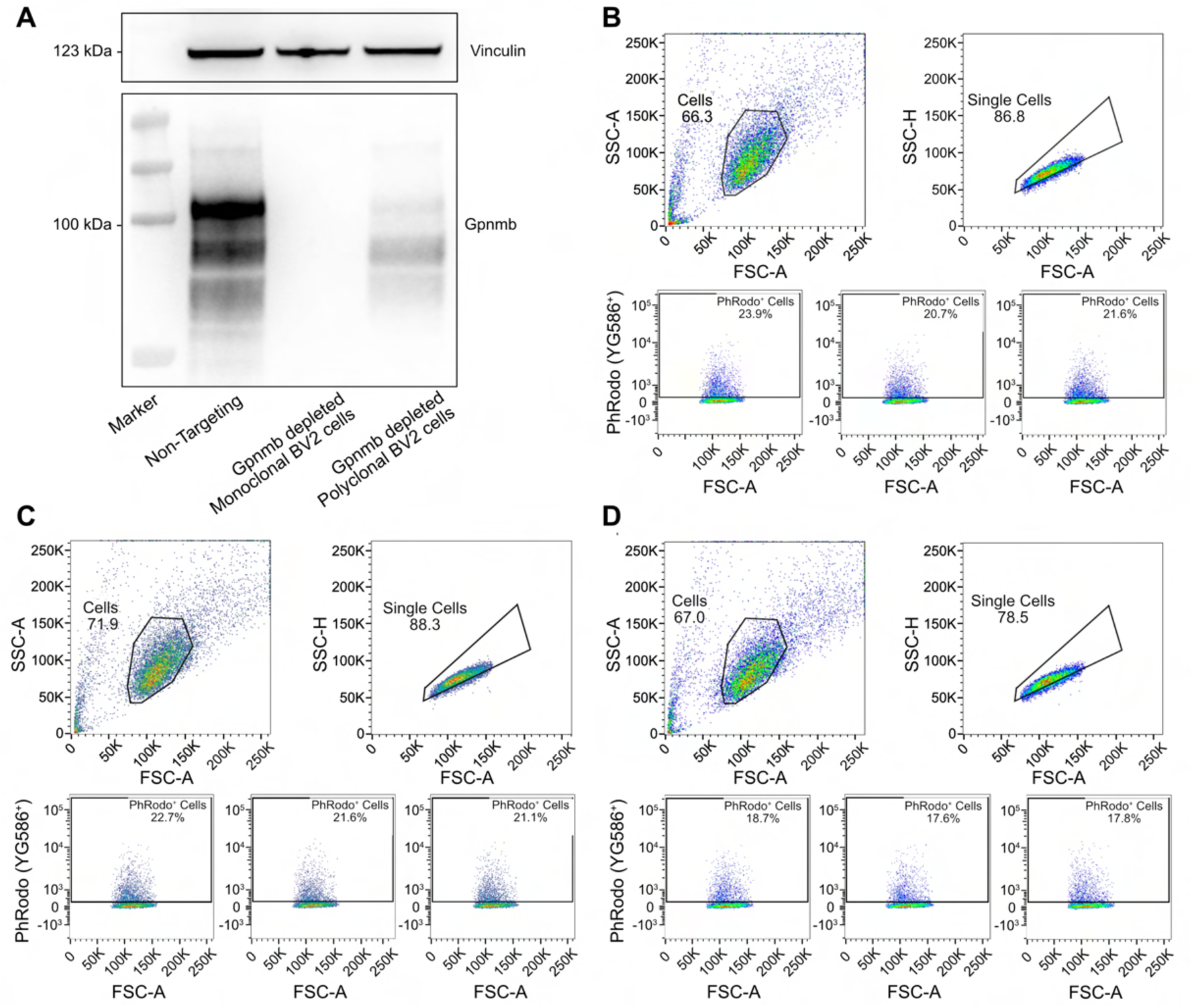
**A)** Western blot showing Gpnmb protein levels in non-targeting control BV2 cells, *Gpnmb*-depleted monoclonal BV2 cells and *Gpnmb*-depleted polyclonal BV2 cells, with Vinculin as a loading control. **B-C)** Top panels: gating strategy for flow cytometry analysis of phagocytic activity in Cas9-expressing BV2 cells. (B) Cells transduced with non-targeting plasmids and (C) cells transduced with a single guide RNA targeting the *Gpnmb* gene, incubated with Phrodo-labeled apoptotic CAD5 cells. Bottom panels: Percentages of Phrodo^+^ cells are indicated from three independent experiments. **D)** Similar to (C), but with an isolated monoclonal BV2 line completely devoid of the *Gpnmb* gene.

**Supplementary Figure 17.**
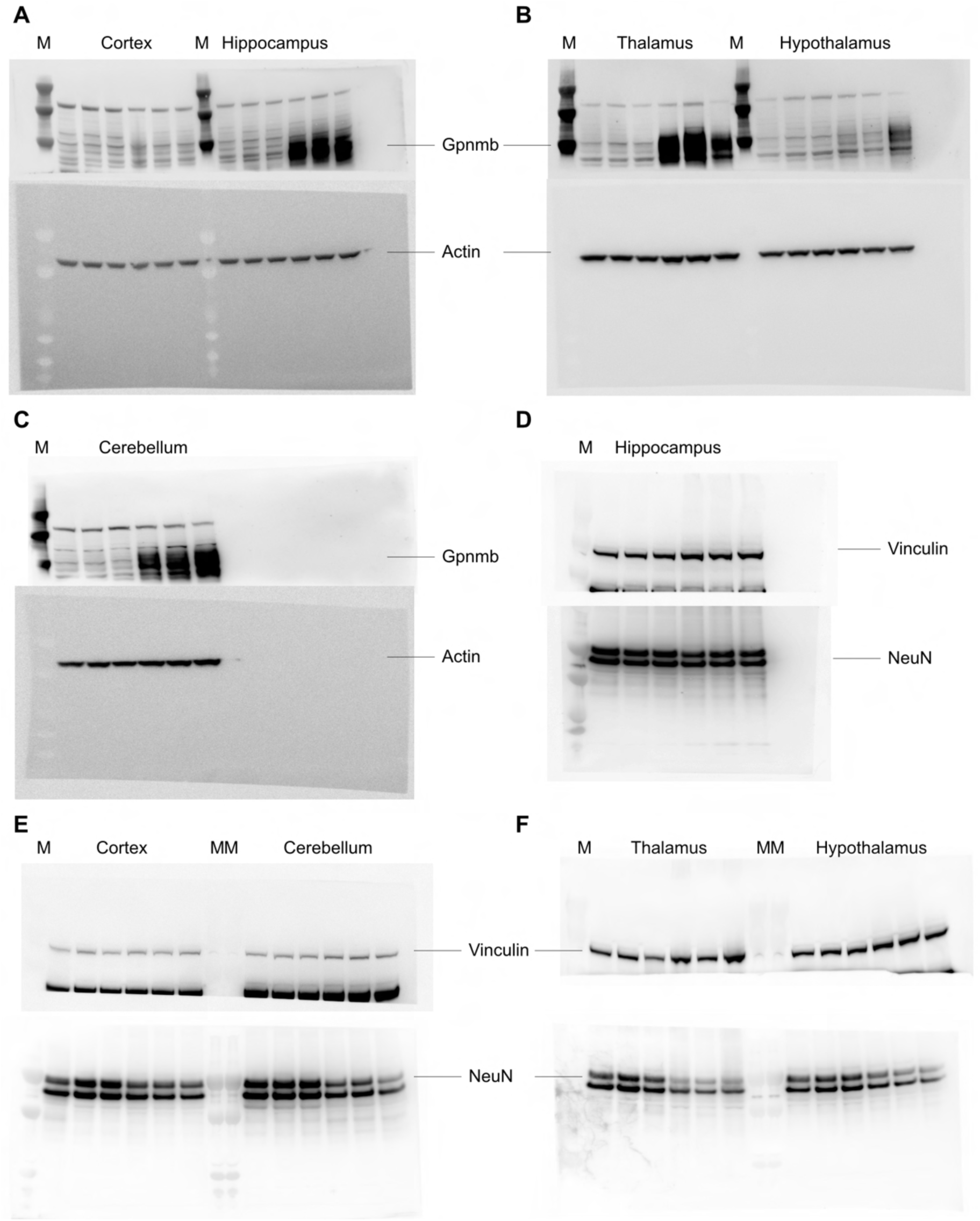
Uncut Western blot of Figure 2B and Figure 5A left panel. Western blots showing Gpnmb and Actin levels in **A**) cortex and hippocampus, **B**) thalamus and hypothalamus and **C**) cerebellum. Western blots of Vinculin and NeuN in **D**) hippocampus, **E**) cortex and cerebellum and **F**) thalamus and hypothalamus. M represents the molecular weight marker. The membranes were cut in two sections and stained with the targeted antibody.

**Supplementary Figure 18.**
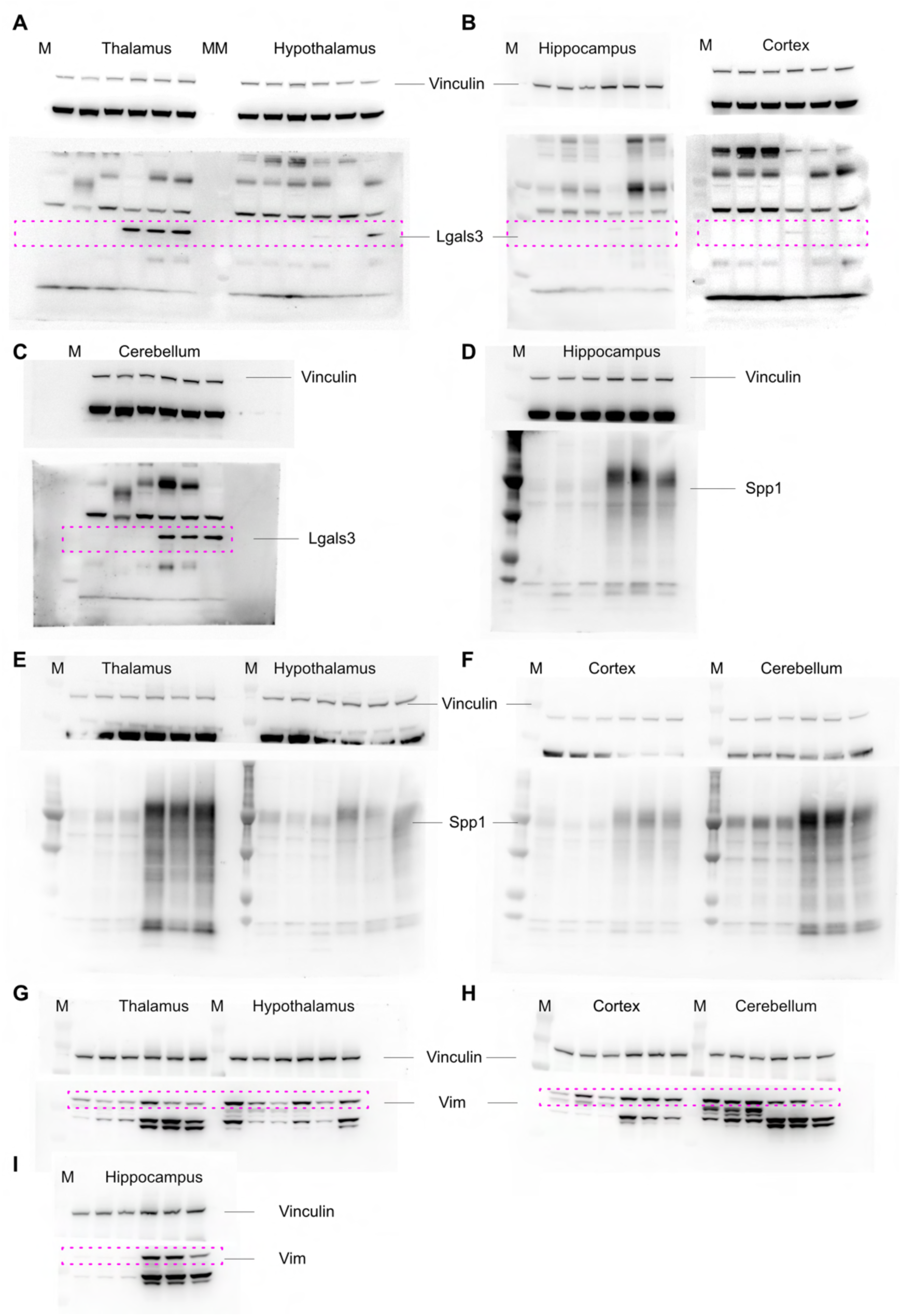
Uncut Western blot of Supplementary Figure 9 A-C. Western blots showing Lgals3 and Vinculin levels in **A**) thalamus and hypothalamus, **B**) hippocampus and cortex and **C**) cerebellum. Western blots of Vinculin and Spp1 in **D**) hippocampus, **E**) thalamus and hypothalamus and **F**) cortex and cerebellum. Western blots of Vinculin and Vim in **G**) thalamus and hypothalamus, **H**) cortex and cerebellum and **I**) hippocampus. Pink-dotted rectangles highlight the bands corresponding to the target antibody among other bands. M represents the molecular weight marker. The membranes were cut in two sections and stained with the targeted antibody.

**Supplementary Figure 19.**
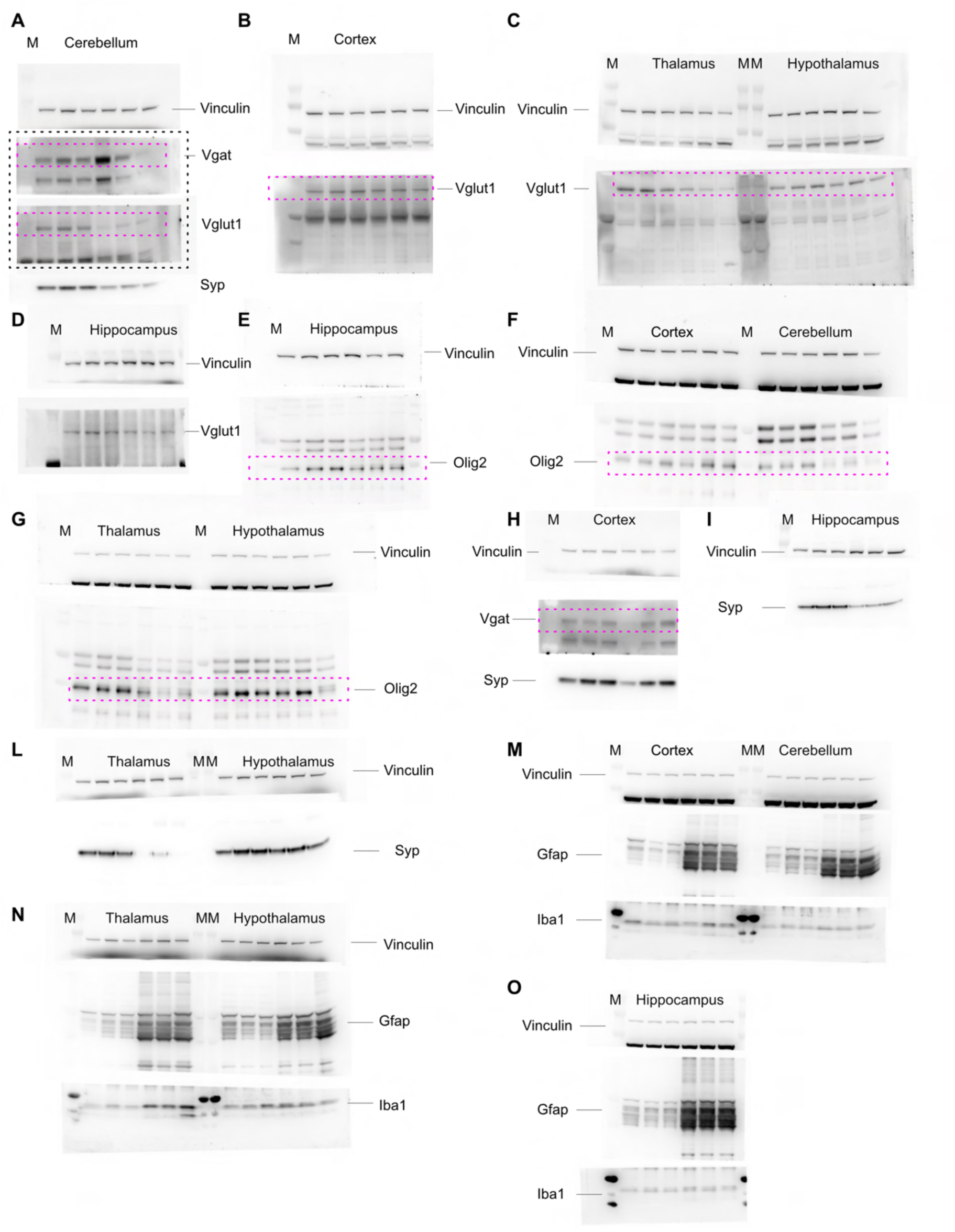
Uncut Western blot of Figure 5A (middle and right panels) and Supplementary Figure 10 A-C. **A**) Western blot was divided into three sections and probed for Vinculin (upper section), Vgat and Vglut1 (middle section), and Syp (lower section) in the cerebellum. The black-dotted rectangle indicates the same blot section, initially stained with the Vglut1 antibody, then stripped and re-probed with Vgat. Western blots of Vinculin and Vglut1 **B**) cortex, **C**) thalamus and hypothalamus, **D**) hippocampus. Western blots of Vinculin and Olig2 in **E**) hippocampus, **F**) cortex and cerebellum and **G**) thalamus and hypothalamus. **H**) Western blots of Vinculin, Vgat and Syp in cortex. Western blot of Vinculin and Syp in **I**) hippocampus, **L**) thalamus and hypothalamus. Western blots of Vinculin, Gfap and Iba1 in **M**) cortex, cerebellum, **N**) thalamus, hypothalamus and **O**) hippocampus. Pink-dotted rectangles highlight the bands corresponding to the target antibody among other bands. M represents the molecular weight marker. The membranes were cut in several sections and stained with the targeted antibody.

**Supplementary Figure 20.**
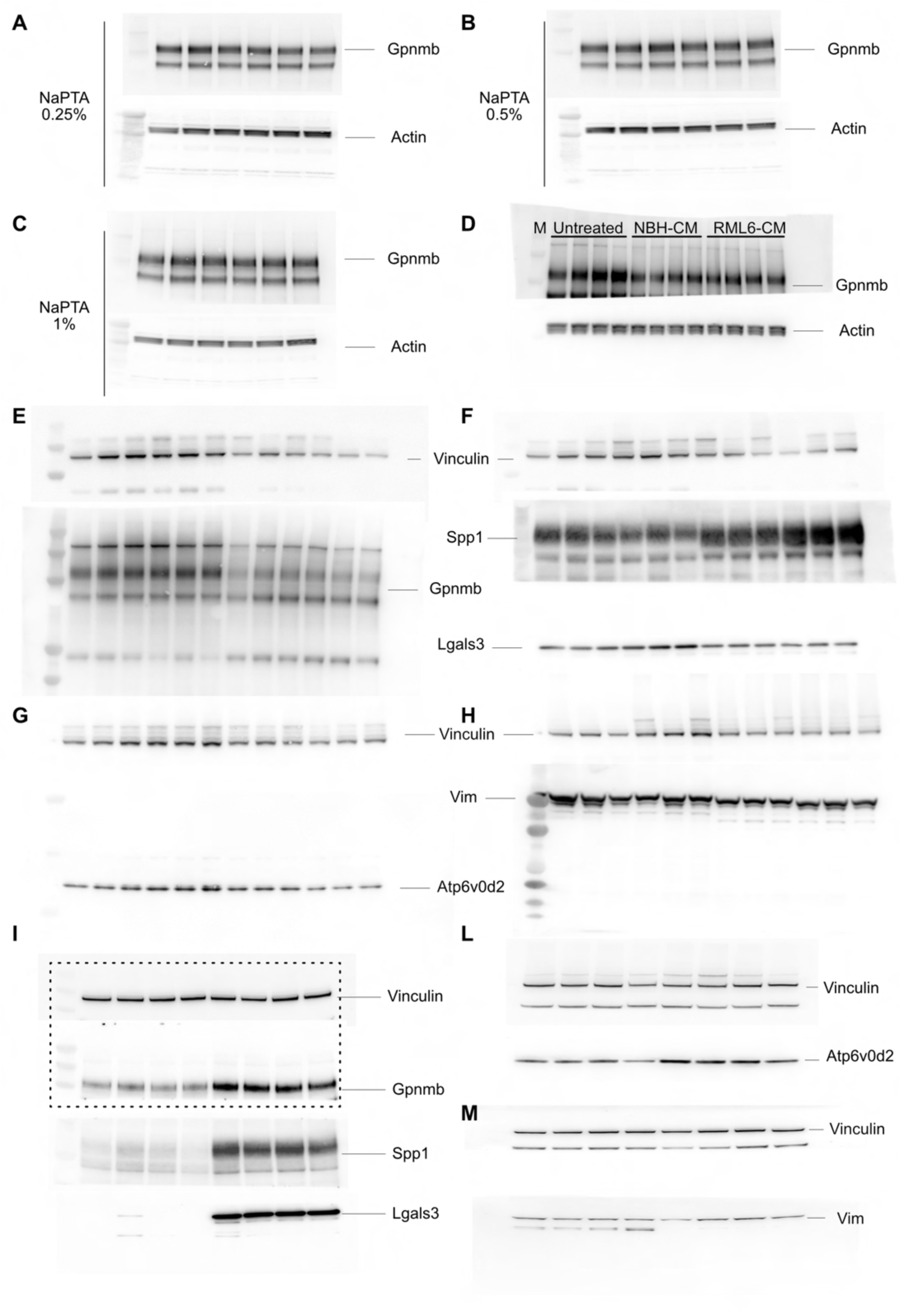
Uncut Western blot analysis corresponding to Supplementary Figure 12C-D, 13A and Figure 6A. Western blots showing Gpnmb and actin in BV2 treated with **A**) 0.25%, **B**) 0.5% and **C**) 1% concentration of NaPTA prion-enriched brain homogenate **D**) Western blot showing Gpnmb and Actin in untreated, NBH-CM, and RML6-CM treated samples. Untreated samples were not included in the manuscript. Western blots showing **E**) Gpnmb and Vinculin as loading control; Gpnmb was first stained, then stripped and re-stained with Vinculin; **F**) Spp1, Lgals3, and Vinculin as the loading control; the membrane was cut in three sections and then stained with targeted antibodies; **G**) Atp6v0d2 and Vinculin as loading control; **H**) Vim and Vinculin as loading control. M represents the molecular weight marker. Uncut Western blots showing **I**) Gpnmb, Spp1, and Lgals3 were analyzed, with Vinculin used as a loading control. The membrane was divided into three sections. The black-dotted rectangle highlights the same section, where the upper portion was first stained for Gpnmb, then stripped and re-probed for Vinculin. Additionally, uncut Western blot showing **L**) Atp6v0d2 and **M**) Vim with related Vinculin as loading control on membranes cut in two sections.

## Notes

### Competing Interest Statement

The authors have declared no competing interest.

### Summary of Updates

This revised version of the manuscript addresses minor errors present in the initial submission and includes new validation experiments. Specifically, we corrected typographical inconsistencies and clarified statements throughout the main text and figure legends. Additionally, we have added a substantial new dataset using a complex human in vitro system to validate our key findings on GPNMB upregulation in microglia. These new experiments confirm that apoptotic stimuli trigger the emergence of a GPNMB-positive microglial phenotype with enhanced phagocytic activity, thereby strengthening the translational relevance of our observations. Minor formatting updates to figures and supplementary material were also implemented.

